# Nuclear lamin isoforms differentially contribute to LINC complex-dependent nucleocytoskeletal coupling and whole cell mechanics

**DOI:** 10.1101/2021.05.12.443683

**Authors:** Amir Vahabikashi, Suganya Sivagurunathan, Fiona Ann Sadsad Nicdao, Yu Long Han, Chan Young Park, Xianrong Wong, Joseph R. Tran, Gregg G. Gundersen, Karen L. Reddy, G.W. Gant Luxton, Ming Guo, Jeffrey J. Fredberg, Yixian Zheng, Stephen A. Adam, Robert D. Goldman

## Abstract

The ability of a cell to regulate its mechanical properties is central to its function. Emerging evidence suggests that interactions between the cell nucleus and cytoskeleton influence cell mechanics through poorly understood mechanisms. Here we show that A- and B-type nuclear lamin isoforms distinctively modulate both nuclear and cellular volume and selectively stabilize the linker of nucleoskeleton and cytoskeleton (LINC) complexes that couple the nucleus to cytoskeletal actin and vimentin. We reveal, further, that loss of each of the four-known lamin isoforms in the mouse embryonic fibroblasts differentially affects cortical and cytoplasmic stiffness as well as cellular contractility, and then propose a LINC complex mediated model that explains these impaired mechanical phenotypes. Finally, we demonstrate that loss of each lamin isoform softens the nucleus in a manner that correlates with loss of heterochromatin. Together, these findings uncover distinctive roles for each lamin isoform in maintaining cellular and nuclear mechanics.

## Introduction

The ability of mammalian cells to regulate their mechanical response to environmental forces is fundamental to their physiological function and motile behavior. Cellular mechanics are facilitated by the complex cytoskeletal networks that extend from the nucleus to the cell periphery. It has also been shown that the nucleus plays a significant role in cellular mechanics, as enucleated cells containing intact intracellular cytoskeletal systems exhibit altered mechanical properties and migratory behavior (1, 2). Taken together, it is clear that the interactions between the nucleus and the cytoskeleton are important for regulating cellular mechanics and motility.

The nuclear envelope (NE) is a specialized organelle that separates the nucleus from the cytoplasm. There is increasing evidence that the NE provides an interface for linking the nucleoplasm and the genome to the various cytoplasmic cytoskeletal systems and the extracellular environment (3). The NE contains a double membrane that is a subdomain of the endoplasmic reticulum (ER), and the nuclear lamina (NL), which is a complex fibrillar meshwork of lamin intermediate filaments and their associated proteins that are closely juxtaposed to the nucleoplasmic face of the inner nuclear membrane. The lamins are type V intermediate filament proteins that are subdivided into either A-types (lamins A (LA) and C (LC)) or B-types (lamins B1 (LB1) and B2 (LB2)). LA and LC are alternatively spliced products of the *LMNA* gene that are expressed in most differentiated cell types, while the ubiquitously expressed LB1 and LB2 proteins are encoded by the *LMNB1* and *LMNB2* genes, respectively (3). Each lamin isoform assembles into a distinct meshwork, but the individual meshworks interact in ways that affect the structure of the other meshworks through unknown mechanisms (4). Underlying and interacting with these meshworks are large domains of heterochromatin called lamina associated domains (LADs) (5).

The NL is physically coupled to the cytoskeleton through a NE-spanning molecular bridge known as the linker of nucleoskeleton and cytoskeleton (LINC) complex (6). Early studies in the mouse embryonic fibroblasts (MEFs) show that deletion of A-type or B-type lamins perturbs the organization of the perinuclear cytoskeleton and significantly softens the cell (7-9). However, the individual contributions from each of the four-known nuclear lamin isoforms to cellular mechanics remain unclear.

Here, we conduct morphometric analyses to show that loss of A- or B-type lamins distinctively affects nuclear and cellular shape and volume. We employ atomic force microscopy (AFM), optical tweezers microrheology (OT), and traction force microscopy (TFM) to reveal that the four lamin isoforms differentially contribute to the stiffness of the cell cortex, the cytoplasm, and the contractile state of MEFs. We then use fluorescence recovery after photobleaching (FRAP) and RNA interference to demonstrate that individual lamin isoforms are central to the stability and function of the LINC complexes that couple the nucleus to the actin and the vimentin intermediate filament (VIF) cytoskeletons. Based on these findings, we propose a novel LINC complex-mediated model that explains the impaired cell mechanics incurred in the absence of each lamin isoform. We further show that loss of each lamin isoform compromises nuclear mechanics and that these resulting changes correlate with changes of the levels of heterochromatin. Together, our studies establish specific roles performed by each lamin isoform in maintaining nuclear and cellular morphology and mechanics.

## Results

### A and B-type lamins affect nuclear and cellular morphology in an opposing manner

Changes in the shape and volume of cells and their nuclei impact cellular mechanical phenotypes (mechanotypes) (10-12). Therefore, we investigated the effects caused by the deletion of specific lamin isoform-encoding genes on nuclear and cellular morphology in MEFs. To do this, we performed confocal imaging of our previously described homozygous lamin knockout (KO) MEF lines (LA/C-, LB1-, and LB2-) as well as their wild type (WT) controls (13). Since recent studies suggest that LA and LC perform distinct roles in establishing the cellular mechanophenotype (14, 15), we also probed the specific contributions of these alternative splice variants in MEFs stably expressing short hairpin RNAs (shRNAs) that knockdown (KD) either endogenous LA (LA KD) or LC (LC KD) (Figs. S1A-B).

Maximum projections and orthogonal sections of nuclei revealed dramatic nuclear shape changes in the lamin KO MEFs as compared to the WT MEFs (Fig. 1A). WT MEF nuclei generally had an oblate spheroid profile with only a few surface distortions, while the profiles of nuclei in the LA/C-MEFs were altered with an increased number of surface irregularities and distortions. In contrast, the LB1- and the LB2-MEF nuclei were less oblate and appeared rounder than the WT MEF nuclei, and also lacked any readily observable distortions (Fig. 1A). To quantify the changes in nuclear shape, we obtained confocal z-sections and analyzed the sphericity of the cells using Imaris software. We found that the nuclei in all the lamin KO or KD MEF lines were significantly rounder than those in the WT MEFs (Figs. 1B and S2A). To test if re-expressing the depleted lamin isoform in its respective KO MEFs rescues nuclear shape, we created MEF lines that stably express LA, LB1, or LB2 in the LA/C-, the LB1-, or the LB2-MEFs respectively (Figs. S1C-D). We were unable to adequately rescue LC expression in LA/C-MEFs for these experiments, so those data are not included in our analyses. We found that the nuclei in all of our rescued KO MEF lines exhibited significant flattening towards the oblate nuclear profile observed in the WT MEFs (Fig. 1B).

**Figure 1.**
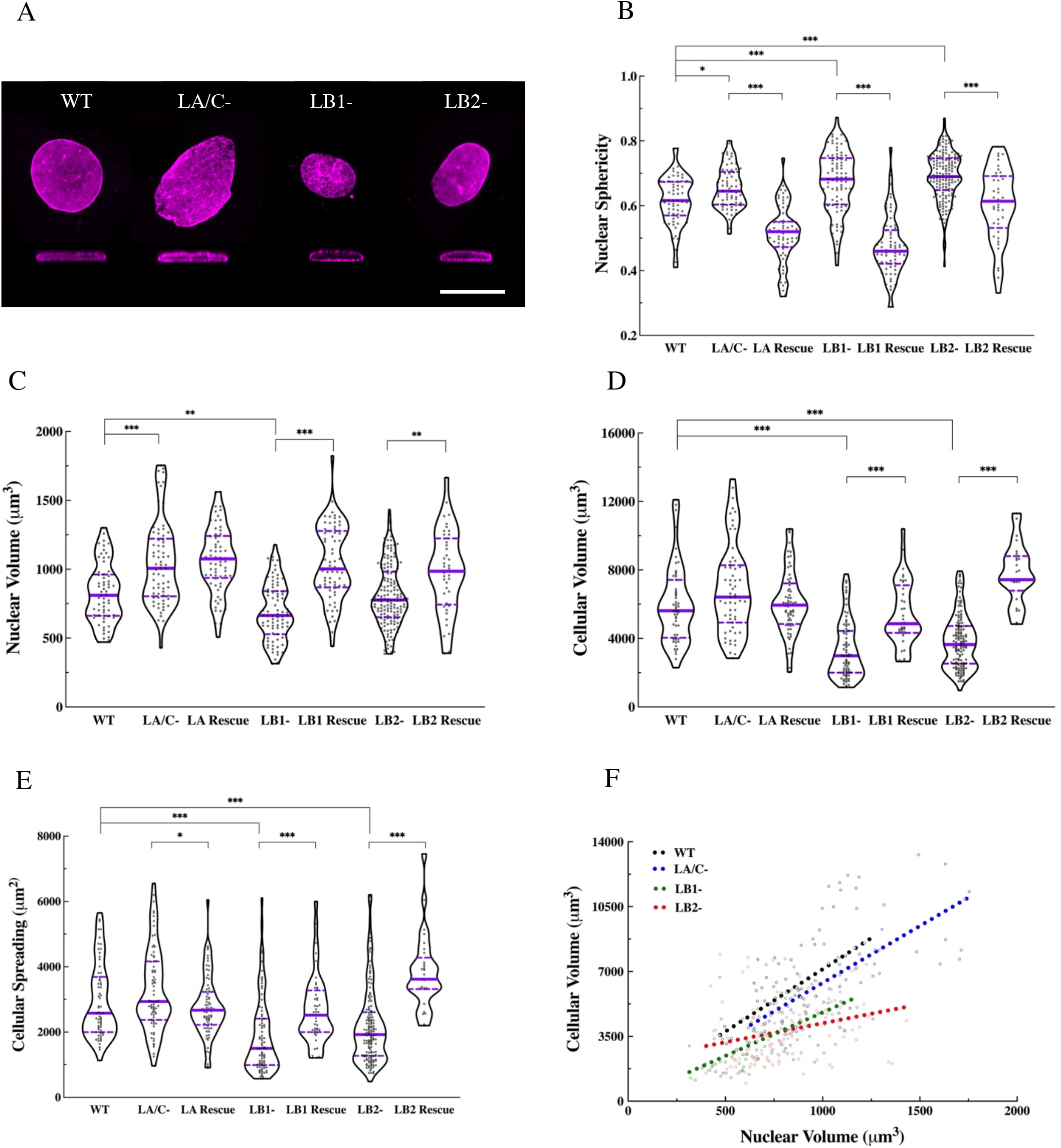
Nuclear lamins are required for the maintenance of nuclear and cellular morphology. **(A)** Representative maximum projections and orthagonal views of the nuclei in the indicated MEF lines stained for the nuclear lamin; The nucleus in the WT, LA/C-, and the LB2-MEF is stained with anti-LB1 and the nucleus in the LB1-MEF is stained with anti-LB2. Violin plots of the average **(B)** nuclear sphericity **(C)** nuclear volume **(D)** cellular volume, and **(E)** cellular spreading area in the WT (n = 3, 55 cells), lamin KO (n = 3, 59-135 cells), and the rescued lamin KO MEFs (n=3, 42-93 cells). **(F)** Scatter plot of cellular vs. nuclear volume in the WT (slope = 6.72, *P <* 0.0001), LA/C-(slope = 6.14, *P <* 0.0001), LB1-(slope = 4.69, *P <* 0.0001), and LB2-(slope = 2.04, *P <* 0.01) MEFs. The regression slopes in LB1- and LB2-MEFs are significnatly different from WT and LA/C-MEFs (*P* < 0.0001). The solid bars in the violin plots represent the median and the dashed lines mark the 25^th^ and 75^th^ percentiles. \**P* < 0.05, \*\**P* < 0.01, \*\*\**P* < 0.001. Scale bar is 20 μm.

Measurements of nuclear volume showed that, relative to the WT MEFs, nuclei were significantly larger in the LA/C-, LA KD, and LC KD MEFs, smaller in the LB1-MEFs and, almost the same size in the LB2-MEFs (Fig. 1C and Fig. S2 B). To determine if LA was sufficient to rescue this phonotype in LA/C-MEFs, we exogenously expressed LA in these cells. LA was not sufficient to restore normal nuclear volume, suggesting a non-redundant role for LC (Fig. 1C). These data are in contrast to the re-expression of LB1 or LB2 in LB1- or LB2-MEFS, respectively, in which a significant increase in nuclear volume was noted. (Fig. 1C).

In addition to the lamin isoform-dependent changes in nuclear shape and volume, we found that cellular volume was also dependent on the lamin isoforms expressed. Both WT and lamin isoform deficient cells were stained for F-actin and subjected to 3D confocal microscopy. Using 3D renderings, we determined that LA/C-MEFs had a slightly larger cellular volume than WT MEFs, whereas the cellular volumes of the LB1-MEFs and the LB2-MEFs were significantly reduced (Fig. 1D). Depletion of LA with shRNA also resulted in a significant increase in the cellular volume of LA KD MEFs, but LC depletion had a small opposite effect on cellular volume in the LC KD MEFs that did not reach statistical significance (Fig. S2C). Re-expressing the missing isoform tended to restore cellular volume to the levels observed in the WT MEFs by slightly decreasing it in the rescued LA/C-MEFs and significantly increasing it in the rescued LB1- and LB2-MEFs (Fig. 1D).

Previous studies suggest a relationship between the extent of cellular spreading and stiffness (11, 16). Therefore, we examined the effect of lamin isoform KO or KD on cell spreading area. Our analyses show that LA/C-MEFs spread slightly more than WT MEFs, while LB1- and LB2-MEFs spread much less (Fig. 1E). Similar to the pattern observed for cellular volume, depletion of LA or LC had contrary effects on cell spreading where LA KD MEFs and LC KD MEFs displayed significantly increased and decreased cell spreading areas, respectively (Fig. S2D). We also found that the cell spreading area of the rescued LA/C-MEFs was significantly decreased approaching WT levels, while both rescued LB1- and LB2-MEFs increased to WT levels or even greater (Fig. 1E).

Since nuclear and cellular volume tend to be positively correlated (17, 18), we next determined the effect of lamin isoform expression on this relationship. Consistent with previous reports (18), linear regression analyses found a significant correlation between the nuclear and cellular volume in WT MEFs, lamin KO MEFs, and lamin KD MEFs (Figs. 1F and S2E). However, the slopes of the regression lines calculated for LB1-, LB2-, and LC KD MEFs were significantly different from those calculated for WT, LA/C-, and LA KD MEFs (Figs. 1F and S2E), suggesting that different modes of nucleocytoskeletal interactions may exist across these cell lines.

The size of the nucleus has also been positively correlated with an increase in cell size during cell cycle progression (19). To determine if the differences in nuclear and cellular size related to lamin isoform expression may be biased by alterations in cell cycle distribution, we quantified the DNA content of each of the MEF lines using flow cytometry. Our analyses show insignificant differences in cell cycle distribution and therefore the differences in nuclear and cellular size of the various MEF lines are not due to the accumulation of cells at a specific stage of the cell cycle (Fig. S3).

### Loss of A- or B-type lamins distinctively compromises cellular mechanics

The finding that MEFs with altered lamin composition exhibit changes in nuclear and cellular morphology suggests that both nuclear and whole cell mechanics are also altered in these cells. To investigate this possibility, we developed a multipronged biophysical approach. Using AFM with large round probes (radius (R) = 5 μm), we characterized the bulk stiffness of the cytoplasm as previously described (20). These measurements show that the cytoplasm in all the lamin KO MEFs is significantly softer relative to the WT MEFs, with LB1- and LB2-MEFs being the softest (Fig. 2A). In addition, depletion of LC significantly decreased the cytoplasmic stiffness while depletion of LA had a negligible effect (Fig. S4A). Re-expressing LA in LA/C-MEFs or LB1 in LB1-MEFs stiffened the cytoplasm to levels observed in the WT MEFs (Fig. 2A). Rescuing the expression of LB2 in LB2-MEFs also increased the cytoplasmic stiffness of these cells, but not to the WT levels (Fig. 2A).

**Figure 2.**
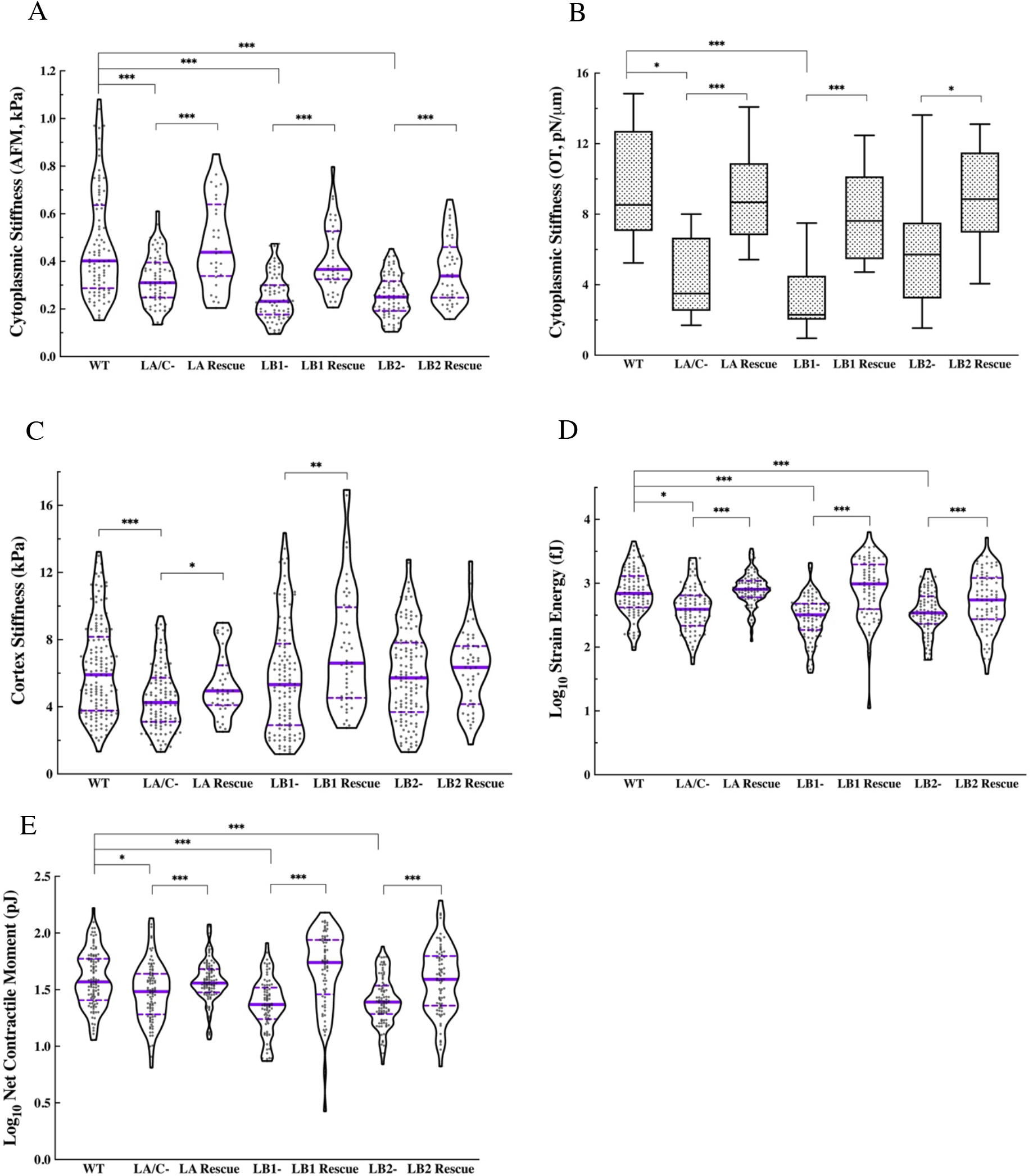
The loss of A- and B-type lamins distinctively influence cellular mechanics. **(A)** Violin plots of the AFM round probe measurements of the cytoplasmic stiffness in the WT (n = 4, 75 cells), lamin KO (n = 4, 69-75 cell), and rescued (n = 4, 30-40 cells) MEFs. **(B)** Box plots of OT measurements for the cytoplasmic stiffness in the WT (n = 1, 9 cells), lamin KO (n = 1, 9-13 cells), and rescued lamin KO (n = 1, 10-12 cells) MEFs. **(C)** Violin plots of the AFM sharp probe measurements for the apical cortical stiffness in the WT (n = 4, 147 cells), lamin KO (n = 4, 117-130 cells), and rescued (n = 4, 44-50 cells) MEFs. **(D)** Violin plots of the logarithmically transformed SE in the WT (n = 2, 105 cells), lamin KO (n = 2, 80-94 cells), and rescued (n = 2, 82-98 cells) MEFs. **(E)** Violin plots of the net contractile moment in the WT (n = 2, 105 cells), lamin KO (n = 2, 81-97 cell), and rescued (n = 3, 82-98 cells) MEFs. The solid bars in the violin plots represent the median and the dashed lines mark the 25th and 75th percentiles. The bars and the whiskers in the box plots represent the median and the minimum/maximum, respectively. *P < 0.05, **P < 0.01, ***P < 0.001.

As a more direct measure of the cytoplasmic stiffness, we used OT-based active microrheology (21). Our OT measurements were consistent with those obtained with the AFM round probe, demonstrating that LA/C- and LB1-MEFs had significantly softer cytoplasm compared to the WT MEFs, and that LB2-MEFs exhibited a lesser reduction in the cytoplasmic stiffness (Fig. 2B). Furthermore, the cytoplasmic stiffness increased to WT levels upon rescuing the expression of the missing lamin isoforms in the corresponding lamin KO MEFs (Fig. 2B). Similar to the AFM measurements, the LC KD MEFs had a significantly softer cytoplasm while LA KD MEFs exhibited a slight decrease in their cytoplasmic stiffness (Fig. S4B).

Cell shape and spreading are known to associate with cortex stiffness (22-24). The relationships between lamin expression and cell spreading (see Fig. 1E) suggest that there may also be changes in the mechanical properties of the cell cortex in MEFs with altered lamin composition. We therefore used our previously established AFM method with a sharp pyramidal probe (R = 20 nm) to measure the apical cortex stiffness of the different MEF lines (20). The results revealed that the cortex of LA/C-MEFs was significantly softer than the cortex of the WT MEFs, while cortical stiffness was mostly unaffected in LB1-, LB2-, LA KD, or LC KD MEFs (Figs. 2C and S4C). We further found that rescuing the LA in LA/C-MEFs and LB1 in LB1-MEFs significantly stiffened the cortex (Fig. 2C), but no significant change in cortical stiffening was observed in LB2-MEFs rescued for the LB2.

The alterations in spreading area and cortex stiffness observed in the lamin KO and KD MEFs further suggested that the contractile forces exerted by these cells at their basal surfaces might also be lamin isoform dependent. To determine whether this were the case, we used TFM in each MEF line to measure the net contractile moment (NCM) exerted by each cell upon its substrate and the strain energy (SE) imparted by each cell to its substrate (25). The data show that SE is notably reduced in all lamin KO and KD MEFs relative to the WT controls (Figs. 2D and S4D). This phenomenon was reversed in rescue experiments as detected by the restoration of the WT levels of SE in the LA/C-, LB1-, and LB2-MEFs rescued for LA, LB1, and LB2 respectively (Fig. 2D). Further analyses of the NCM measurements showed a strong agreement with the results from the SE measurements demonstrating significant reduction of the contractile moment in LC KD MEFs as well as all lamin KO MEFs relative to the WT controls, which was restored to the WT levels upon rescuing the expression of the missing lamin isoform (Figs. 2E and S4E).

### Loss of A- or B-type lamins compromises nuclear stiffness and alters heterochromatin levels

Lamins are proposed to regulate nuclear mechanics not only through their non-linear mechanical properties as intermediate filament polymers but also through their influence on the organizational state of chromatin (e.g., euchromatin vs. heterochromatin) (26-29). However, the individual contribution of each lamin isoform to nuclear stiffness is incompletely understood. To begin to address this knowledge gap, we measured nuclear stiffness in the lamin KO, KD, and rescued MEF lines. We measured the bulk stiffness of the nucleus using AFM with large round tips (R = 5 μm) to carry out nano-indentations over the center of the nucleus as described previously (30, 31). The results demonstrate that the nuclei in all lamin KO MEFs were significantly softer than the nuclei in the WT MEFs, with the degree of softening being comparable between the lamin KO lines (Fig. 3A). However, we observed that nuclear stiffness did not change in either LA KD or LC KD MEFs relative to the WT MEFs (Fig. S5A). Measurements performed in the rescued LA/C-MEFs demonstrated that re-expressing LA in the LA/C-MEFs dramatically stiffened the nucleus to levels that were even greater than those observed in the WT MEFs (Fig. 3A). Nuclear stiffness in the rescued LB1-MEFs was restored to the same level as was detected in WT MEFs. However, this was not the case for the rescued LB2-MEFs, where the nucleus became slightly stiffer, but not to the same extent as observed in the WT MEFs (Fig. 3A).

**Figure 3.**
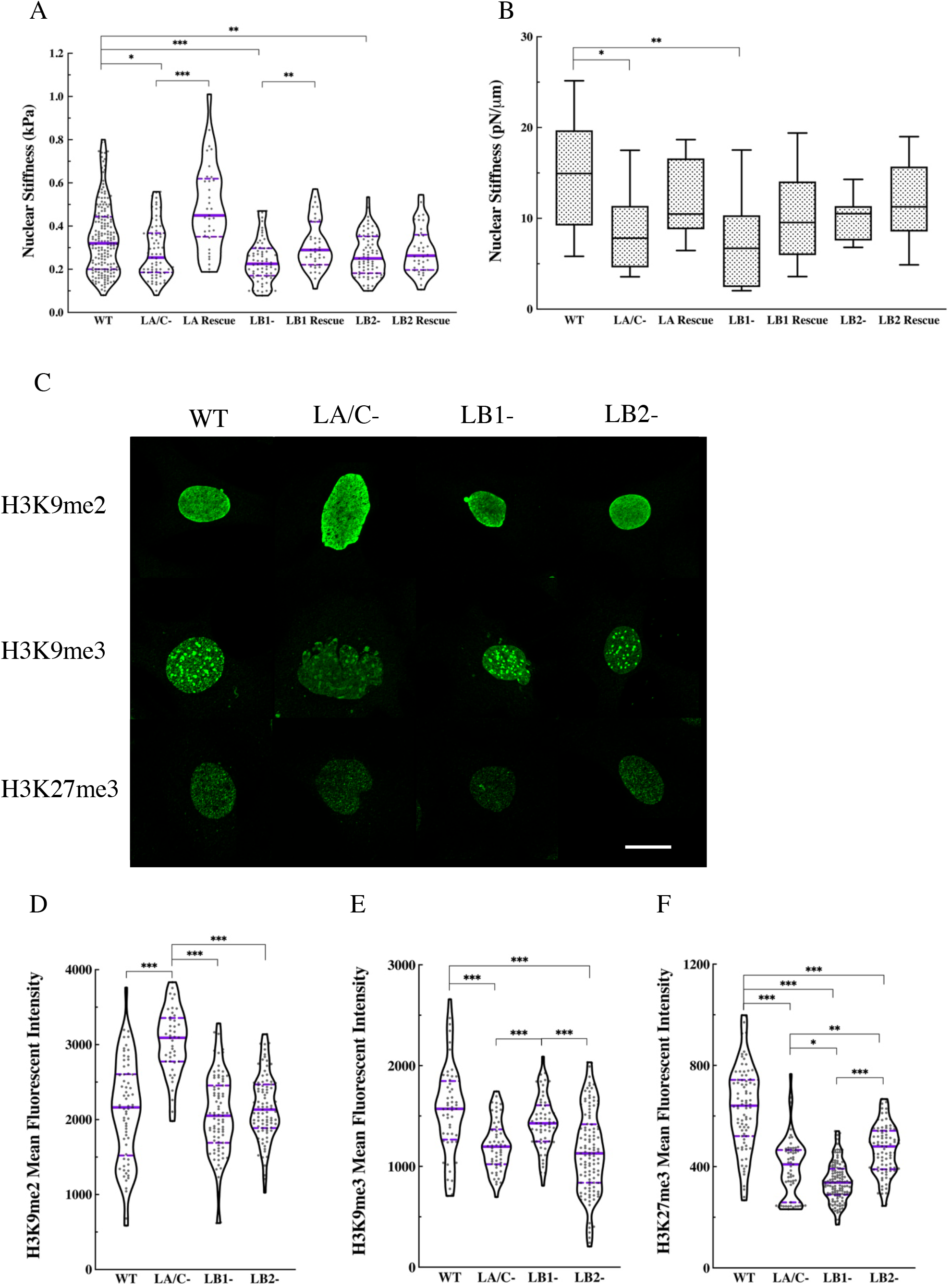
The loss of nuclear lamin isoforms compromises nuclear stiffness and alters heterochromatin levels in MEFs. **(A)** Violin plots of AFM round probe measurements of the nuclear stiffness in the WT (n = 4, 188 cells), lamin KO (n = 4, 69-73 cells), and rescued (n = 2, 33-39 cells) MEFs. **(B)** Box plots of the OT measurements for the nuclear stiffness in the WT (n = 1, 9 cells), lamin KO (n = 1, 9-12 cells), and rescued (n = 1, 9-12 cells) MEFs. **(C)** Representative maximum projections of confocal z-stacks of nuclei in the WT, LA/C-, LB1-, and LB2-MEFs stained for H3K9me2, H3K9me3, or H3K27me3. Violin plots of the mean fluorescent intensity of **(D)** H3K9me2, **(E)** H3K9me3, and **(F)** H3K27me3 in the WT (n = 2, 54-93 cells), LA/C-(n = 2, 44-59 cells), LB1-(n = 2, 61-132 cells), and LB2-(n = 2, 91-117 cells) MEFs. The solid bars in the violin plots represent the median and the dashed lines mark the 25^th^ and 75^th^ percentiles. The bars and the whiskers in the box plots represent the median and the minimum/maximum, respectively. \**P* < 0.05, \*\**P* < 0.01, \*\*\**P* < 0.001. Scale bar is 20 μm.

We also used OT to more directly measure nuclear stiffness in the MEF lines. To achieve this, we located endocytosed latex beads (R = 0.25 μm) in the cytoplasm adjacent to the nucleus and dragged them towards the nucleus to indent the nuclear surface (32). Similar to our AFM results, the OT measurements showed a significant decrease in nuclear stiffness in LA/C- and LB1-MEFs relative to the WT MEFs. In addition, a notable, yet statistically insignificant, decrease in nuclear stiffness was measured in LB2-MEFs compared to the WT MEFs (Fig. 3B). Nuclear stiffness measurements performed in all rescued lamin KO MEFs revealed a notable, but statistically insignificant, increase in nuclear stiffness relative to the WT MEFs (Fig. 3B). In contrast to our AFM studies, the OT measurements indicated that LA KD significantly softened the nucleus, whereas LC KD had a marginal effect on the nuclear stiffness (Fig. S5B).

Small mechanical deformations of the nucleus have been shown to be regulated by the heterochromatin (26). Given the small deformations (< 1 μm) inherent in the AFM and OT assays, we speculated that the observed decreases in nuclear stiffness in the lamin KO and KD MEFs may be, at least in part, related to alterations in the levels of their heterochromatin. This possibility is supported by the findings that changes in the levels of heterochromatic markers (i.e. H3K9me2, H3K9me3, and H3K27me3) are associated with altered nuclear stiffness (27, 33). Therefore, we determined the levels of heterochromatin in lamin KO and KD MEFs by quantitative immunofluorescence using antibodies directed against histone marks for heterochromatin (Figs. 3C-F and S5C-F). Quantification of the mean fluorescence intensity for H3K9me2 showed a significant increase in LA/C-MEFs relative to the WT controls, but the fluorescence intensity levels in LB1-, LB2-, LA KD, and LC KD MEFs remained unchanged (Figs. 3D and S5D). In addition, the mean fluorescence intensities of H3K9me3 were significantly reduced in LA/C- and LB2-MEFs, as compared to the WT and LB1-MEFs (Fig. 3E.). Furthermore, the mean fluorescence intensity of H3K9me3 was also significantly reduced in LC KD MEFs relative to the WT and LA KD MEFs (Fig. S5E).

The mean fluorescence intensity of H3K27me3, was dramatically reduced in all lamin KO MEFs relative to the WT MEFs (Fig. 3F). However, the extent of this decrease was different between the lamin KO MEFs with LB1-MEFs showing significantly lower intensity compared to the LA/C-, LB2-MEFs, and LA/C-MEFs demonstrating significantly lower intensity relative to the LB2-MEFs. LA KD MEFs exhibited increased H3K27me3 intensity relative to the WT MEFs, while the fluorescence intensity in LC KD MEFs was similar to the WT MEFs (Fig. S5F). To further verify the results from the immunofluorescence studies, we examined the expression levels of H3K9me and H3K27me in the WT, lamin KO, and lamin KD MEFs through western blot analysis (Fig. S5G). Data from the western blots show good agreement with the immunofluorescence results, suggesting an overall decrease in the level of these heterochromatic markers in the lamin KO and KD MEFs. These results show that the loss of each lamin isoform reduces heterochromatin levels, which in turn may contribute to the nuclear softening.

### A- and B-type lamins distinctively regulate the dynamics of LINC complexes that bind F-actin and vimentin

We reasoned that the compromised cytoplasmic and cortical stiffness of the A- or B-type lamin KO MEFs might be related to defects in their ability to form proper nucleocytoskeletal connections via LINC complexes. Given the essential roles of the F-actin and VIF cytoskeletal systems in regulating cortical and cytoplasmic mechanics (20, 34, 35), we hypothesized that the assembly of LINC complexes that bind these cytoskeletal filaments might be defective in lamin KO MEFs.

To test this hypothesis, we first expressed EGFP-tagged mini-nesprin-2 giant (a previously characterized functional nesprin-2G construct (36)), nesprin-3α, SUN1, or SUN2 in the WT and lamin KO MEF lines. Nesprin-2G directly interacts with F-actin (36, 37), while nesprin-3α binds VIFs via plectin (38). Nesprin-2G and nesprin-3α can both interact with SUN1 or SUN2 within the perinuclear space of the NE (39-41). Consistent with previous studies (42, 43), these EGFP-tagged proteins all localized to the NE in WT as well as lamin KO MEFs (Fig. S6A).

In order to determine whether the dynamic properties of the LINC complex proteins were disrupted when lamin isoform expression was altered, we carried out quantitative fluorescence recovery after photobleaching (FRAP) experiments in the NEs of the MEF lines that individually express each of the EGFP-tagged LINC complex proteins and determined the halftime (t_1/2_) of recovery (Fig. S6B). The fluorescence of EGFP-SUN1 and EGFP-SUN2 recovered in the NE bleached zone in all lamin KO MEFs (Fig. 4A-B). However, the normalized t_1/2_ for both EGFP-SUN1 and EGFP-SUN2 was significantly faster in LA/C- and the LB2-MEFs relative to LB1- and WT MEFs (Fig. 4E). Furthermore, the fluorescence of EGFP-SUN2 in LA/C-MEFs tended to recover significantly faster than in the NE of LB2-MEFs (Fig. 4E). Therefore, the mobility of SUN1 and SUN2 is significantly increased in LA/C- and LB2-MEFs as compared to the LB1- and WT MEFs.

**Figure 4.**
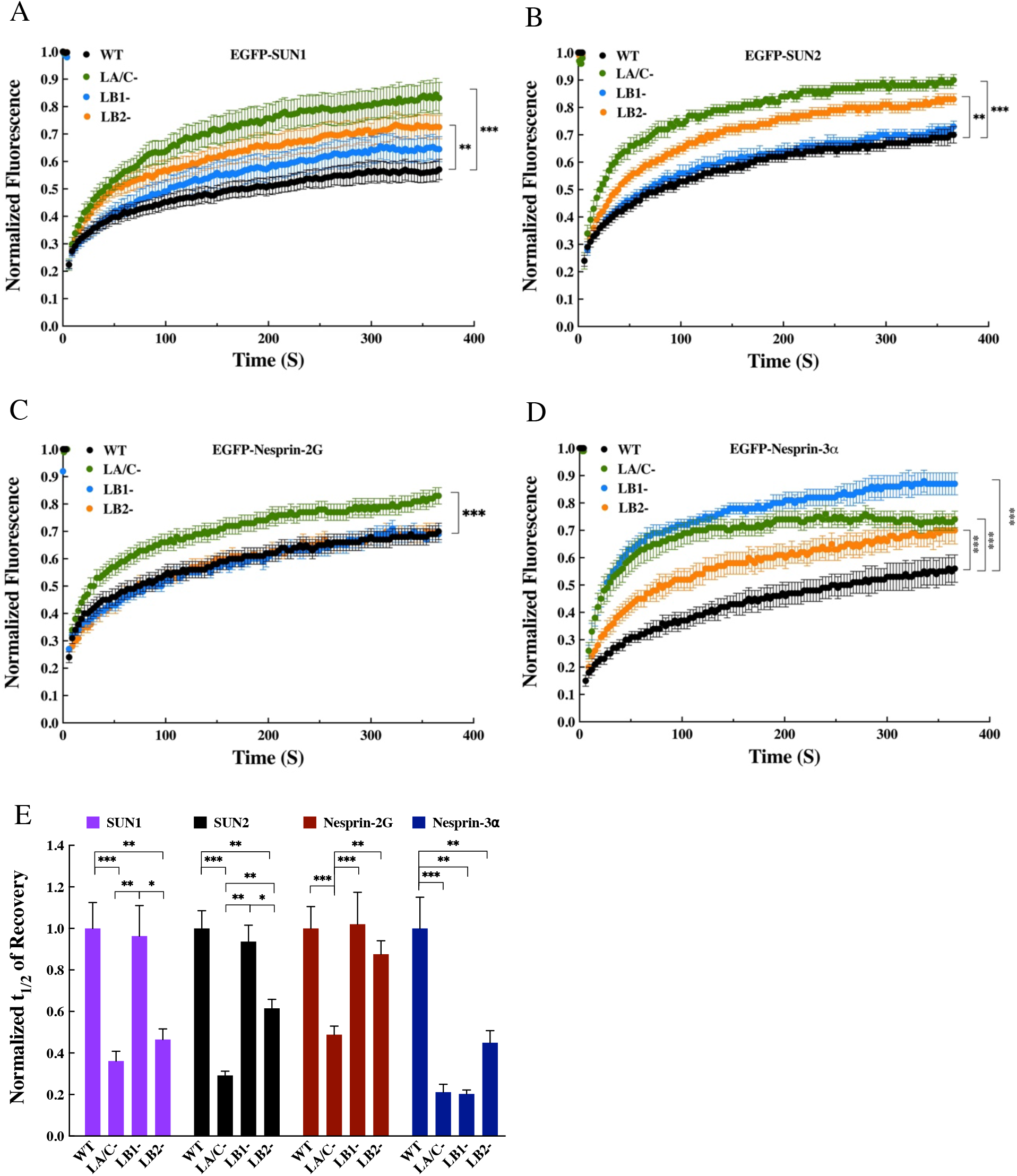
A- and B-type lamins distinctively regulate the dynamics of F-actin or vimentin binding LINC complex components at the NEs of the MEFs. Plots of the normalized fluorescence recovery of **(A)** EGFP-SUN1, **(B)** EGFP-SUN2, **(C)** EGFP-mini-Nesprin-2G, and **(D)** EGFP-Nesprin-3*α* in the NEs of the WT, LA/C-, LB1-, and LB2-MEF lines. **(E)** Bar plots of the normalized t_1/2_ of NE FRAP for the indicated EGFP-tagged LINC complex constructs expressed in the WT, LA/C-, LB1-, and LB2-MEF lines. The reported values are normalized to those obtained in the WT MEFs. See Fig. S6B for the absolute numbers of t_1/2_. (n ≥ 2; 10-15 cells per experimental condition). The data are shown as mean ± SE. \**P* < 0.05, \*\**P* < 0.01, \*\*\**P* < 0.001.

FRAP analyses of EGFP-mini-nesprin-2G revealed a significantly faster recovery in the NE of LA/C-MEFs (Fig. 4C). However, the fluorescence recovery of EGFP-mini-nesprin-2G in the NE was not distinguishable among LB1-, LB2-, and WT MEFs (Fig. 4E). We further observed a dramatic increase in the normalized fluorescence intensity of EGFP-nesprin-3*α* in all lamin KO MEFs (Fig. 4D), which manifests itself in a significantly faster normalized t_1/2_ for EGFP-nesprin-3*α* in the NE of these cells (Fig. 4E). The data also suggest a relatively faster fluorescence recovery in the LA/C- and LB1-MEFs compared to the LB2-MEFs, although this difference is not statistically significant. These findings suggest that the loss of lamins increases the mobility of the F-actin- and VIF-binding LINC complexes within the NEs of the lamin KO MEFs. These observations are also consistent with previous studies that show increased mobility of SUN1, SUN2, and Nesprin-2G in the LA/C-MEFs (42, 44, 45). Taken together, the FRAP studies establish substantial and distinct roles for the A- and the B-type lamins in regulating the mobility and, by extension, the assembly of LINC complexes that couple the nucleus to the actin or vimentin cytoskeletons.

### Inhibition or depletion of the LINC complex recapitulates the mechanical defects caused by lamin knock-out

The FRAP results suggest relationships between the lamins, cellular stiffness, and the ability of cells to assemble functional LINC complexes. Therefore, we hypothesized that inhibition of the function of LINC complex components should result in cellular mechanotypes similar to those observed in the lamin KO MEFs. To test this hypothesis, we used AFM to measure cortical and cytoplasmic stiffness in MEFs expressing one of the three established EGFP-tagged dominant negative LINC complex constructs. These constructs are targeted to the lumen of the ER and the contiguous perinuclear space of the NE by a signal sequence (SS) fused to the N-terminus of EGFP (SS-EGFP). Two dominant negative constructs consist of SS-EGFP fused to the N-terminus of the SUN protein-binding KASH peptides of nesprin-2 or −4 (SS-EGFP-KASH2 or - KASH4) (46), while the third construct consists of SS-EGFP fused to the N-terminus of the KASH peptide-binding luminal domain of SUN1 (SS-EGFP-SUN1^LD^) (46). As a control for SS-EGFP-KASH2 and -KASH4, we used a SS-EGFP-KASH2 construct lacking the four conserved C-terminal amino acids that are critical for SUN protein-binding (SS-EGFP-KASH2^ΔPPPT^) (47). To control for the SS-EGFP-SUN1^LD^ construct, we used a SS-EGFP construct that contains an ER retention sequence at its C-terminus (SS-EGFP-KDEL) (46). We also used non-transfected WT MEFs as a second control for these dominant negative construct expression experiments. Consistent with their ability to inhibit LINC complex assembly (48, 49), immunofluorescence demonstrated that the expression of each of the three dominant negative constructs in WT MEFs displaced endogenous nesprin-2G from the NE to the endoplasmic reticulum (Fig. S7A).

The results of AFM round tip measurements demonstrate that the expression of SS-EGFP-KASH2 or SS-EGFP-KASH4 significantly softened the cytoplasm of WT MEFs when compared to the non-transfected MEFs or MEFs expressing the SS-EGFP-KASH2^ΔPPPT^ control construct (Fig. 5A). Similarly, the expression of SS-EGFP-SUN1^LD^ resulted in a significantly softer cytoplasm in WT MEFs when compared to non-transfected WT MEFs or those expressing the SS-EGFP-KDEL control construct (Fig. 5A). Cortical stiffness measurements with AFM sharp tip probes showed that the expression of SS-EGFP-KASH2 results in a significant softening of the cell cortex in WT MEFs compared to non-transfected controls and a notable, yet statistically insignificant, softening compared to the WT MEFs expressing the SS-EGFP-KASH2^ΔPPPT^ control construct (Fig. 5B). The cortex was also significantly softer in WT MEFs expressing either SS-EGFP-KASH4 or SS-EGFP-SUN1^LD^, as compared to non-transfected WT MEFs or those expressing the appropriate corresponding controls (Fig. 5B).

**Figure 5.**
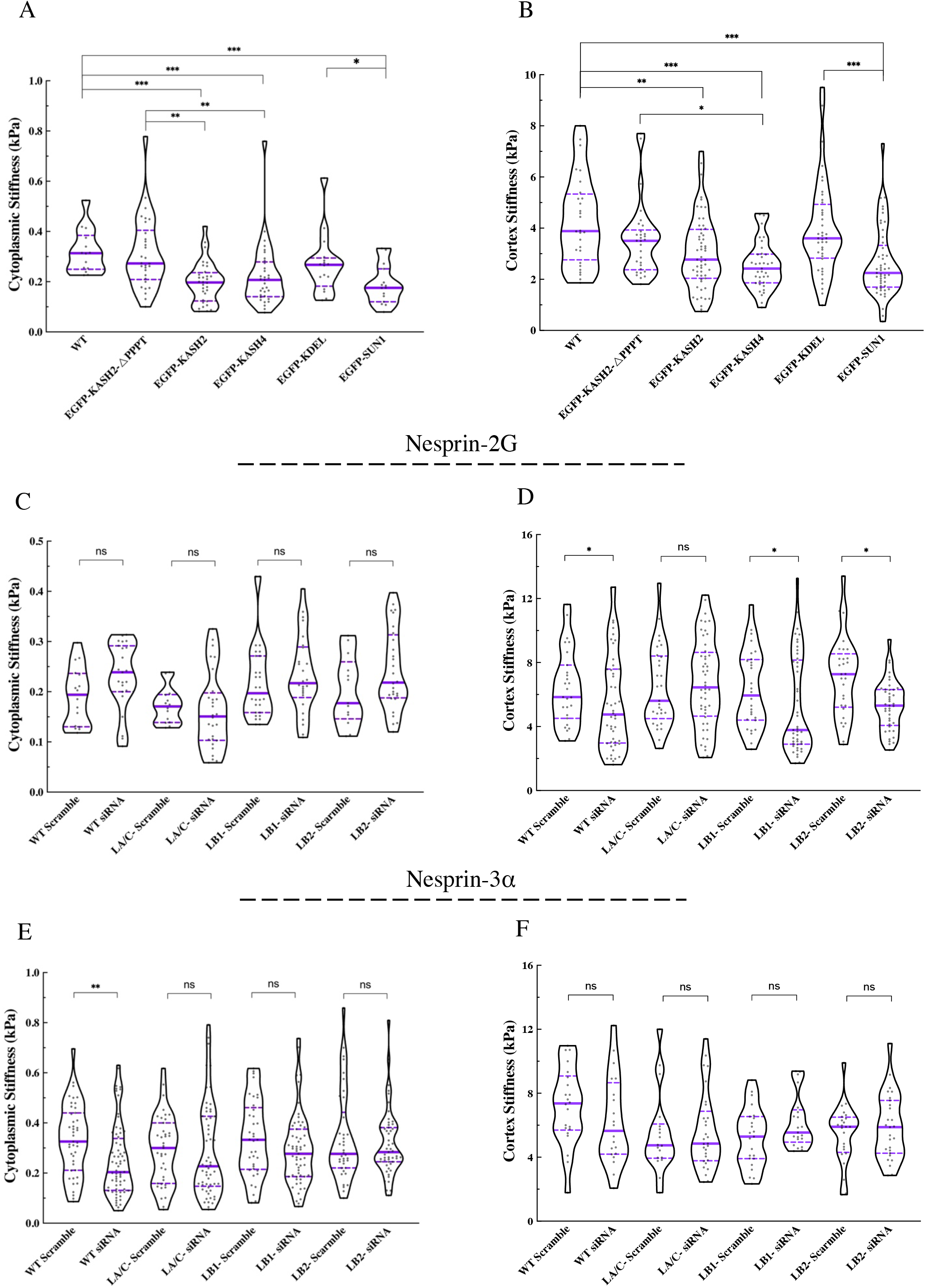
LINC complex inhibition and depletion of nesprin-2G or nesprin-3α recapitulates the cellular mechanotypes caused by the absence of the nuclear lamins in MEFs. Violin plots of the **(A)** cytoplasmic and **(B)** cortical stiffness measurements in non-transfected WT MEFs and upon the expression of the indicated SS-EGFP-tagged dominant negative LINC complex inhibitor constructs or their corresponding controls (At least 15 cells were measured per experiment). Violin plots of the **(C)** cytoplasmic and **(D)** cortical stiffness measurements in the WT, LA/C-, LB1-, and LB2-MEFs treated with either the non-coding control or the nesprin-2 siRNA (n=2, at least 15 cells were measured per experiment). Violin plots of the **(E)** cytoplasmic and **(F)** cortical stiffness measurements in the WT, LA/C-, LB1-, and LB2-MEFs treated with either the non-coding control or the nesprin-3α siRNA (n=2, at least 15 cells per experiment). The solid bars in the violin plots represent the median and the dashed lines mark the 25^th^ and 75^th^ percentiles. The bars and the whiskers in the box plots represent the median and the minimum/maximum, respectively. \**P* < 0.05, \*\**P* < 0.01, \*\*\**P* < 0.001.

Having established a role for the LINC complex in regulating both cortical and cytoplasmic stiffness in WT MEFs, we next reasoned that if lamins were regulators of cellular mechanics through their physical interactions with LINC complexes, then depletion of LINC complex components that bind the F-actin or VIF cytoskeletons should have minimal or no additional effects on the mechanics of the lamin KO MEFs, since the nuclear-cytoskeletal coupling would be impaired at the site of attachment to the lamina in these cells. To test this possibility, we performed AFM measurements in WT and lamin KO MEFs treated with small interfering RNAs (siRNAs) to deplete either endogenous nesprin-2G or nesprin-3α (Figs. 5C-F and S7B). For controls, we performed AFM measurements in WT and lamin KO MEFs treated with scrambled siRNAs. We found that the depletion of nesprin-2G had a slight effect on the stiffness of the cytoplasm in WT MEFs but none of the changes were statistically different from those measured in WT MEFs treated with the control siRNAs (Fig. 5C). However, nesprin-2G-depletion significantly softened the cortex in WT, LB1-, and LB2-MEFs, but had a negligible effect on cortical stiffness of LA/C-MEFs (Fig. 5D). We further found that the depletion of nesprin-3α from WT MEFs resulted in a significantly softer cytoplasm, but it did not change cytoplasmic stiffness in lamin KO MEFs (Fig. 5E). In addition, cortical stiffness was slightly reduced in nesprin-3α-depleted WT MEFs and no detectable changes in cortical stiffness were observed in any of the nesprin-3α-depleted lamin KO MEFs (Fig. 5F). Taken together, these results demonstrate that the A- and B-type nuclear lamin isoforms distinctively interact with F-actin and VIF binding LINC complexes to regulate the mechanical properties of the cell cortex and cytoplasm.

### Loss of A- and B-type lamin isoforms alters cell motility and enhances the levels of double-stranded DNA damage caused by constricted cell migration

Loss or altered expression levels of A- and B-type lamins have been associated with changes in cell migration (7, 50-54). Therefore, we studied the relationship between the altered cellular mechanics observed in lamin KO and KD MEFs and their motility in 2D and 3D-like environments. In a 2D wound healing assay (Fig. S8A), LA/C-MEFs migrated at marginally higher rates than WT MEFs, whereas LB1- and LB2-MEFs moved at significantly higher rates (Fig. 6A). In addition, we found that LA KD MEFs and LC KD MEFs migrated slower and faster than WT MEFs, respectively (Fig. S8B). To examine the relationships between lamin isoforms and cell migration in 3D-like microenvironments, we used transwell migration assays. Based on our nuclear volume measurements (Figs. 1C and S2B), we calculated an average effective radius of 5.4-6.2 μm for the nuclei across all KO and KD MEFs. Since the nucleus is the largest organelle inside the cell (55), we chose to use transwell filters with either 3 or 5 μm pore diameters to induce two levels of constricted migration and thus, mechanical stress. When migrating through the 5 μm pores, LA/C- and LB2-MEFs, but not LB1-MEFs, migrated at significantly higher levels relative to WT MEFs (Fig. 6B). The overall migration of these MEF lines through the 3 μm pores was generally reduced, but the average migration ability was significantly increased in all lamin KO MEFs when compared with the WT MEFs (Fig. 6B). The migration of LA KD and LC KD MEFs across both 3 and 5 μm pore diameters was comparable to that observed with WT MEFs (Fig. S8C). To examine the role of nuclear size in cell migration through the pores, we compared our migration data to the average diameter of the nuclei normalized by the pore diameter (Figs. S8D-E). The results suggest that despite comparable ratios for nuclear to pore diameter, cell migration rates were significantly higher in lamin KO MEFs relative to the WT MEFs.

**Figure 6.**
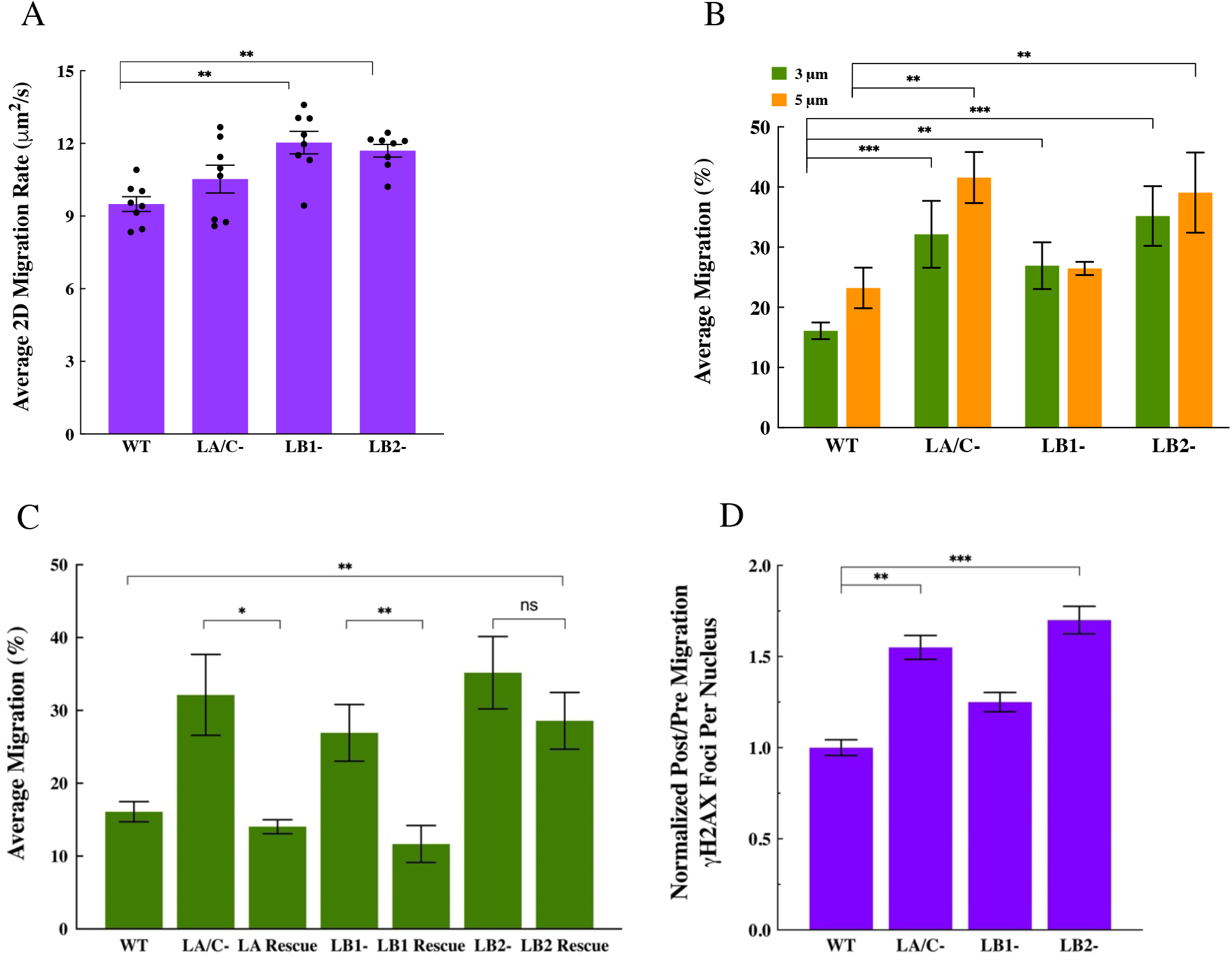
The loss of A- or B-type lamins enhances cell migration and increases constricted migration-induced DNA damage. **(A)** Bar Plots of the quantification of 2D migration rate in a wound healing assay for the WT, LA/C-, LB1-, and LB2-MEFs (n = 8). **(B)** Bar plots of the average migration percentage for the WT, LA/C-, LB1-, and LB2-MEFs through transwell membranes with 3 μm or 5 μm pore diameters. (n ≥ 5 per experiment condition). **(C)** Bar plots of the average migration percentage through transwell membranes with 3 μm pores for the indicated MEF lines (n ≥ 3). **(D)** Bar plots of the normalized post/pre migration γ-H2AX foci counts in the WT, LA/C-, LB1-, and LB2-MEFs after migrating through transwells with 3 μm pores. (n =3, at least 150 cells per experiment condition). Data are shown as mean ± SE. \**P* < 0.05, \*\**P* < 0.01, \*\*\**P* < 0.001.

We next attempted to rescue the migratory behavior of the lamin KO MEFs to the levels exhibited by the WT MEFs by exogenously expressing the missing lamins in these cells and examining their ability to migrate through 3 μm pores. We found that the ability of rescued LA/C- and rescued LB1-MEFs to migrate through 3 μm pores was indistinguishable from the WT MEFs. In contrast, the migration of rescued LB2-MEFs was slightly decreased relative to the WT MEFs (Fig. 6C). We also observed that for all examined MEF lines, the average nuclear area of the cells that migrated through the pores was smaller than those that remained at the top of the transwell (Figs. S8F-G). This observation is consistent with previous studies that demonstrate smaller pores select for cells with smaller nuclei (18, 56).

Cell migration through constricted spaces is associated with an accumulation of damaged DNA within the nucleus (18, 57-59). Therefore, we determined if the loss of lamin isoforms affects migration-induced DNA damage in lamin KO and KD MEFs by immunostaining the cells on transwell membranes for γ-H2AX (Fig. S8H), a marker of double-stranded DNA breaks (60). The normalized counts for γ-H2AX foci per nucleus was then compared between cells that migrated through the transwell pores and those that did not. These results showed that LA/C- and LB2-MEFs exhibited over 50% and 70% increase in the number of γ-H2AX foci as compared to the WT MEFs, respectively (Fig. 6D). DNA damage was also increased by over 20% in LB1-, LA KD, and LC KD MEFs relative to the WT MEFs, but these increases were not statistically significant (Figs. 6D and S8I).

## Discussion

Our results demonstrate that each of the four major lamin isoforms contributes to mammalian cell mechanics in a distinctive LINC complex-mediated manner. In particular, we find that A-type lamins engage with nesprin-2G- and nesprin-3α containing LINC complexes and therefore F-actin and VIFs, to respectively modulate the cortex and cytoplasmic stiffness of the MEFs, while B-type lamins harness nesprin-3α, and subsequently VIFs, to regulate cytoplasmic stiffness. We further show that all lamin isoforms contribute to the contractile state of MEFs. These findings extend the functional reach of the nuclear lamins well beyond their established roles as key determinants of nuclear stiffness to more global regulators of both the mechanical and contractile properties of the whole cell.

We employed both AFM nanoindentation analyses and OT microrheology measurements to characterize cytoplasmic mechanics related to each lamin isoform. Our previous studies using finite element modeling demonstrated that spherical AFM tips (R = 5 μm) measure the bulk modulus of the cytoplasm because of the large size of the indenting probe, whereas OT uses the motion of a much smaller bead (R = 0.25 μm) to locally characterize the stiffness of the cytoplasm (20, 34, 61). Here, we show that the cytoplasm is much softer in the LA/C-, LB1-, the LB2-MEFs relative to the WT MEFs. Early studies used active and passive microrheology to show that the cytoplasm in LA/C-MEFs is significantly softer when compared to the WT MEFs (7-9). Our results confirm the former observations made in LA/C-MEFs and expand those studies to reveal that the cytoplasm is also notably softer in both LB1- and LB2-MEFs. These findings are consistent with reported parallel microfiltration measurements that show increased deformability of LA/C- and LB1-MEFs relative to WT MEFs (62). Using specific silencing vectors, we have also shown that depletion of endogenous LC but not LA compromises cytoplasmic stiffness in MEFs. This result is further supported by the finding that LC expression levels correlate with the mechanical properties of whole cells (15). The strong agreement between our OT and AFM round probe measurements suggests that the loss of specific lamin isoforms softens the cytoplasm at both local and global scales. We hypothesize that this effect could be due to dysfunctional nucleocytoskeletal connections with the VIF cytoskeleton, since it is a key determinant of the cytoplasmic stiffness (20, 21, 34, 63). Our FRAP results for EGFP-nesprin-3α support this hypothesis by revealing a highly destabilized nuclear connection to the VIFs via LINC complexes. This is also consistent with previous reports on altered organization of perinuclear VIFs in LA/C-MEFs (64). Furthermore, our nesprin-3-depletion experiments show that disrupting LINC complexes that couple VIFs to the nucleus significantly soften the cytoplasm in WT MEFs but has a minimal effect in all lamin KO MEFs. These findings along with the data from the rescue experiments suggest that the meshworks of LA/C, LB1, and LB2 interact, directly or indirectly, with LINC complexes to mediate the coupling of the nucleus to the VIF cytoskeleton. Surprisingly, the removal of any single lamin isoform is sufficient to disrupt this connection and consequently, compromise cytoplasmic stiffness.

Our AFM sharp tip measurements demonstrate that the loss of LA/C, but not LB1 or LB2, significantly softens the cell cortex in MEFs, demonstrating the importance of understanding the different roles played by each lamin isoform in regulating the mechanical properties of the entire cell. Cortex stiffness depends on actomyosin driven cortical tension (35), the extent of connectivity within the actomyosin network (23), and its interactions with the VIF cytoskeleton, as we and others have recently reported (20, 65). A recent study showed that the depletion of LA/C does not change phosphorylation of myosin light chain, which is a well-established marker of contractility (66). However, other studies performed in LA/C-MEFs have documented impaired anchorage of linear arrays of nesprin-2G and SUN2-containing LINC complexes known as transmembrane actin-associated nuclear (TAN) lines during rearward nuclear movement in cells polarizing for directional migration (44), loss or disruption of the highly contractile perinuclear actin caps found on the dorsal nuclear surface (67), and deregulated actomyosin remodeling (66). These reports of the altered dynamics of the actomyosin cytoskeleton are in agreement with ours and other’s (42) observations of the increased mobility of the primary F-actin-binding LINC complex protein nesprin-2G in the NE of LA/C-MEFs. Interestingly, we found that the loss of B-type lamins does not change the mobility of nesprin-2G in the NE and has a minimal effect on the stiffness of the apical cell cortex. These results suggest that nesprin-2G-mediated coupling of the actomyosin cytoskeleton to the dorsal surface of the nucleus potentially modulates the stiffness of the apical cortex through an interaction with LA/C. This proposed model is strengthened by the results of our siRNA experiments, where we show that nesprin-2G-depletion dramatically softens the cortex in WT, LB1-, and LB2-MEFs. However, depletion of the nesprin-2G had almost a negligible effect on cortical stiffness in LA/C-MEFs. We further show that the depletion of either LA or LC has no effect on cortical stiffness and that restoring the expression of LA in LA/C-MEFs is sufficient for significant cortex stiffening. These findings suggest that either LA or LC is likely sufficient to properly immobilize nesprin-2G in the NE. Another finding of our study is that while the loss of LB1 in LB1-MEFs resulted in a small, insignificant drop in cortical stiffness, re-expressing LB1 in these cells significantly stiffened their cortex. We believe this may likely be due to the previously shown effect of LB1 expression on the LA/C meshwork, where LB1-MEFs, but not LB2-MEFs, showed a substantially enlarged LA/C mesh size relative to the LA/C meshwork in WT MEFs (4). In other words, it is likely that in addition to the composition of the nuclear lamina, the organization of its meshwork could likely modulate cytoskeletal dynamics and cell stiffness.

Our studies further reveal a correlation between the reduced contractility in lamin KO and KD MEFs and the disrupted LINC complex dynamics within their NEs. These observations are consistent with recent studies that show that the magnitude of cellular traction forces is associated with an interactive feedback between nucleocytoskeletal connectivity, cytoskeletal tension, and focal adhesions (1, 68). The restoration of contractility to the WT level upon re-expression of the missing lamin isoforms in the corresponding lamin KO MEFs further emphasizes the central role of intact nucleocytoskeletal connections during the regulation of cellular contractility. In particular, it is possible that nesprin-2G-mediated coupling of the nucleus and the actomyosin cytoskeleton might influence cellular contractility (68). However, an interesting finding of our studies is that both LB1- and LB2-MEFs exhibit significantly reduced cellular contractility, which is restored upon re-expressing of their respective missing lamin isoform. These observations suggest that nesprin-3α-mediated nucleocytoskeletal connections and VIFs are likely significant contributors to cellular contractility and the generation of cellular traction forces as well. This proposition is consistent with previous studies that show VIFs governing the alignment of actin-based traction forces (69) and that the loss of VIFs significantly reduces cellular contractility in MEFs by over 30% (20).

The loss of lamins is associated with altered nuclear shape and mechanics (52, 70). Originally, LA/C was proposed as the dominant lamin isoform in regulating nucleus stiffness (70). However, similar to our findings, there is emerging evidence showing that both LB1 and LB2 also contribute to nuclear stiffness (28, 54). The AFM and OT indentations made in these studies cause small deformations in the nucleus (< 1μm), a deformation regime that is dominantly determined by the heterochromatin (26). Consistent with this notion, we find that the significant softening of nuclei observed in the lamin KO MEFs correlates with their notably reduced levels of H3K9me3 and H3K27me3 heterochromatic markers. These observations are in agreement with previous studies that have shown softening of the nucleus upon reduction of the levels of the heterochromatin markers H3K9me3 (33) and H3K9me27 (27).

The results of studies reported here show an increased migratory behavior of lamin KO MEFs through confined (5 μm pore) or strictly confined (3 μm pore) microenvironments. This is consistent with other studies that suggest nuclear stiffness is a physical limit for confined cell migration (71, 72). Further validation of this proposition comes from the clear correlation between the extent of enhanced nuclear stiffness in the rescued lamin KO MEFs and their reduced migration through 3μm pores. Moreover, the enhanced confined migration of both the A- and B-type lamin KO MEFs is accompanied by an increase in the levels of confined migration-induced double-stranded DNA damage. These results are in agreement with previous reports of the increased deformation and double-stranded DNA damage observed in cells with softer nuclei post-migration (50, 73).

In summary, our findings implicate the four-known nuclear lamin isoforms as key determinants of whole cell mechanics. We show that the loss of A-type lamins compromises cortical and cytoplasmic stiffness as well as cellular contractility, while the lack of B-type lamins affects cytoplasmic stiffness and cellular contractility. We reveal that these distinct mechanical changes correlate with the selective interactions of the A- and B-type lamins with the LINC complexes containing nesprin-2G and/or nesprin-3α, which physically couple the nucleus to the actomyosin and VIF cytoskeletons, respectively. We further demonstrate that loss of each lamin isoform reduces nuclear heterochromatin levels, softens the nucleus, and promotes cell migration in constricted microenvironments resulting in significantly increased levels of double-stranded DNA damage. These insights expand our understanding of several fundamental nuclear lamina-dependent cellular processes, such as mechanotransduction and cell migration as well as the pathological conditions associated with mutations in nuclear lamins, including cancer and the nuclear laminopathies (e.g. dilated cardiomyopathy, muscular dystrophy, and the accelerated aging disorder Hutchinson-Gilford progeria syndrome) (74-76).

## Materials and methods

### Cell culture

WT and lamin KO MEFs were isolated from their corresponding mouse embryos as described previously (77) and immortalized by transduction with a retrovirus expressing SV40 T antigen from pBABE-puro SV40LT (pBABE-puro SV40LT was a gift from Thomas Roberts (Addgene plasmid # 13970; http://n2t.net/addgene:13970; RRID:Addgene_13970)). All cells were cultured in DMEM medium containing sodium pyruvate (Life Technologies; Grand Island, NY) and supplemented with 10% fetal bovine serum (Atlanta Biologicals, MN) and 1% penicillin/streptomycin (Milipore Sigma, MO) at 37°C with 5% CO_2_.

### Immunofluorescence and microscopy

WT, KO and KD MEFs were seeded on glass coverslips (#1.5). For all studies, MEFs were fixed in 4% paraformaldehyde for 10 min at RT followed by permeabilization in PBS-containing 0.05% Triton X-100 for 10 min at RT. Following fixation and permeabilization, MEFs were incubated in 10% normal goat or donkey serum containing primary antibodies for 30 min at RT. Subsequently the coverslips were washed 3X (3 min each) in PBS containing 0.01% Tween followed by a final wash in PBS alone. Cells were next incubated with the appropriate secondary antibodies for 30 min at RT and then washed in PBS as above. For morphometric studies, the glass cover slips were mounted using a glycerol mounting media with p-phenylenediamine anti-fading agent. For all other studies, coverslips and membranes were mounted on glass slides using Prolong Diamond (Thermo Fisher Scientific, MA). The following primary antibodies were used: rabbit anti-LA (1:500; #323; (78)), mouse monoclonal anti-LC (1:50; EM-11, Novus Biologicals), goat polyclonal anti-LB1 (1:500; SC-6217, Santa Cruz Biotechnology), rabbit monoclonal anti-LB2 (1:100; EPR9701, Abcam), mouse anti-ƔH2AX (1:500; EMD Millipore), rabbit anti-H3K9me2 (1:1000; #39239, Active Motif), rabbit anti-H3K9me3 (1:1000; ab8898, Abcam), rabbit anti-H3K27me3 (1:1,000; 07-449, Millipore Sigma), rabbit anti-nesprin-2G (36), rabbit anti-nesprin-3 (kind gift from Dan Starr (79)), and Hoechst 33342 (1:10,000). The secondary antibodies used in our study were AlexaFluor 488- or 647-labeled donkey anti-rabbit (1:400; Thermo Fisher), AlexaFluor 488-, 568-, or 647-labeled donkey anti-mouse (1:400; Thermo Fisher), donkey anti-goat Alexa Fluor 488, 568 (1:400; Thermo Fisher), and Alexa Fluor PLUS 488- or 568-labeled goat anti-rabbit (1:400, Thermo Fisher). The actin cytoskeleton was labeled with Alexa Fluor 488- or 568-labeled phalloidin (1:400; Thermo Fisher). All the imaging experiments presented in this work were conducted using a Nikon A1R confocal scope equipped with 20x (air, 0.3 NA), 40x (oil, 1.2 NA), and 60x (oil, 1.4 NA) objectives.

For morphometric analyses, cells were labeled with lamin isoform staining and Alexa Fluor 488 phalloidin to demarcate the boundaries of the nucleus and the cell cortex respectively. The nuclei in the WT, LA/C-, and the LB2-MEFs were stained with anti-lamin B1, while the nuclei in the LB1-MEFs were stained with anti-Lamin B2. Confocal z-stacks (200 nm optical sections) of the nucleus and the whole cell were acquired using a 60x (oil, 1.4 NA) objective. For nuclear surface area/volume measurements and cell volume measurements, confocal z-stacks were imported to Imaris to create 3D renderings as described below. For the cell spreading analyses, maximum projections of the images of actin staining were analyzed using ImageJ (National Institutes of Health), and the cell periphery was manually traced to measure the cell spreading area.

For chromatin immunofluorescence, fluorescence intensity measurements were made using confocal z-stacks (200 nm optical sections) of nuclei using a 60x (oil, 1.4 NA) objective. Maximum projections of the images of nuclei were obtained in ImageJ using automatic threshold detection. The background fluorescence was determined by selecting a 50 × 50 pixel cell free area. The average fluorescence intensity of the nuclei was determined, and the average background intensity present within the image was subtracted (27). All imaging and threshold settings were kept constant when generating and processing the data for comparison of the different MEF lines.

### 3D measurements and sphericity analysis

Confocal z-stacks (200nm optical sections) of the cell/nucleus from WT (n = 63), lamin KO (n = 72-159), lamin KD (n = 72-98), and rescued (n = 42-93) MEFs were imported into Imaris v9.6 (Oxford Instruments, MA) to generate 3D renderings as described previously (80). Cell and nucleus area/volumes were calculated using 3D renderings. Nuclear sphericity is defined as 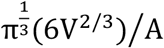, where V is the nuclear volume and A is the surface area. A value of 1 corresponds to a perfect sphere with the value reducing to less than 1 as the object becomes more oblate.

### Transfection and RNA interference

The siRNA pool (3 gene specific siRNAs, SYNE2 Mouse siRNA Oligo Duplex (Locus ID 319565), Cat# SR423729) and the scrambled negative control (Trilencer-27 Universal Scrambled Negative Control siRNA Duplex, Cat# SR30004) for Nesprin-2 were purchased form ORIGENE. Nesprin-3 siRNA SMARTpool (4 individual siRNAs, Mouse Syne3 siRNA, Gene ID: 212073) and its ON-TARGETplus Non-targeting control pool (Cat# D-001810-10-05) were from Horizon Discovery. All transfections were performed using Lipofectamine 3000 (Thermo Fisher, MA) following the manufacturer’s recommended protocol. siRNA transfection on day one at 20nM for Nesprin-2 and 50nM for Nesprin-3 followed by a second transfection at similar siRNA concentration after 24hrs gave the highest knockdown efficiency in the examined MEFs 48hrs after initial transfection as evidenced by immunofluorescence examination for Nesprin-2 and Nesprin-3. We thus adopted this method for our siRNA studies.

Generation of the LA KD and LC KD MEFs. Selective KD of LA or LC was carried out by transduction of WT MEFs with lentiviruses prepared from pLKO.1 constructs containing a neomycin/G418 resistance cassette as well as the shRNAs targeting the non-conserved regions of LA or the 3’ UTR of the LC mRNA (81). Viral particles were produced by 293FT cells that were co-transfected with the LA- or LC-targeting pLKO.1 constructs as well as psPAX2 and pVSV-G. Viral particles were harvested from these cells 48 hrs following transfection. The transduced MEFs were then selected with 400µg/ml G418 for 7 days and subsequently screened for LA or LC KD by immunofluorescence microscopy and immunoblotting with antibodies specific for either LA (rabbit #321) or LC (EM-11 Novus Biologicals).

### Generation of rescued lamin KO MEFs

The re-expression of lamin isoforms in the KO MEFs was achieved by transiently transfecting these cells with pPyCAGIP constructs that encode mouse LA, LB1, or LB2 using Lipofectamine 3000 (Thermo Fisher Scientific) (82). Forty-eight hours following transfection, the cells were selected for 2 days with 2 µg/ml puromycin and immediately used for AFM. The expression of lamins was confirmed by immunofluorescence with antibodies specific for each lamin isoform as described in the section on Immunofluorescence and microscopy. The average lamin concentration in the rescued lamin KO MEFs was determined by quantitative immunoblotting with the following antibodies: LA (rabbit 266, 1:500 (83)); LB1 (Proteintech AG3631, 1:2000); LB2 (mouse 2B2 1:1000;(83)). The average amount of specific lamin expression in each of the rescued MEF lines compared to WT MEFs was 63% for LA, 80% for LB1, and 290% for LB2 determined as described in the section on Immunoblotting.

### Cell cycle analysis

Cells were harvested and washed twice with PBS at 335xg for 5 min followed by fixation in 70 % ethanol for 30 min at −20°C. After fixation, the cells were diluted with PBS and centrifuged at 1126xg for 10 min followed by a PBS wash. The fixed cells were then stained by resuspending in 0.5 ml of DAPI solution (1 µg/ml of DAPI in 0.1 % Triton X-100) for 30 min at 4°C. The stained cells were analyzed on a BD FACS Melody Cell Sorter (BD Biosciences, CA).

### Immunoblotting

Whole cell lysates of MEFs were prepared by trypsinizing cells, quenching in DMEM with 10% FBS, washing twice in PBS by centrifugation at 200xg and finally solubilizing the pellet in 1x SDS sample buffer containing equal numbers of the WT MEFs, the lamin KO MEFs, and the lamin KD MEFs. Lysates were run in triplicate on an SDS-PAGE gel and electrophoretically transferred to nitrocellulose. The membranes were stained for total protein with the Revert Protein Staining kit (LI-COR) and imaged at 700 nm using a LI-COR Odyssey Fc. The protein stain was then removed using the destaining solution included in the kit. After blocking the membranes in 5% non-fat dry milk in PBS with 0.1% Tween 20, the blots were probed with specific antibodies in blocking buffer overnight at 4°C. Antibodies used: LA/C rabbit 266 (1:500) (83); LB1 rabbit Proteintech AG3631 (1:2000); LB2 mouse 2B2 (1:1000) (83); H3K9me mouse Cell Signaling 6F12 (1:2000); H3K27me rabbit Cell Signaling C36B11 (1:1000). After washing 3X for 5 mins each in PBS with 0.1% Tween 20, the blots were probed with 1:15,000 dilution of IRDye 800CW Donkey anti-mouse IgG or 800CW donkey anti-rabbit secondaries (LI-COR) in 5% non-fat dry milk in PBS with 0.2% Tween 20 for 45 mins at RT. After washing twice for 5 mins each in PBS containing 0.2% Tween 20, the blot was allowed to air-dry in the dark. Imaging of the membranes was performed on a LI-COR Odyssey Fc and the resulting images were analyzed using LI-COR Empiria software. The amount of protein present in each lane was quantified from the Revert protein stain image and used to correct for sample loading across the different lanes of the gel.

### Transwell migration assays

Polycarbonate transwell membranes with pore diameters of either 3 μm or 5 μm (Corning, NY) were used. Membranes were pre-coated with rat-tail collagen 1 (50 μg/mL, Thermo Fisher, MA) for 30 min, rinsed with PBS, incubated with pre-warmed growth medium for 60 min, and then seeded with 10,000 cells/well followed by incubation at 37°C for 18 hours. The membranes were then fixed and stained for immunofluorescence with LA, LB1, and ƔH2AX antibodies. Cells were gently removed from either the top or bottom of the membrane with a cotton swab and the membrane was then mounted on glass coverslips using Prolong Diamond (Thermo Fisher Scientific). The membranes were imaged with a 20x (air, 0.3 NA) objective in 600 × 600 μm^2^ fields of view (10 consistent locations per condition) and nuclear count and area were determined by lamin staining and through thresholding and automated particle analysis in ImageJ (84). Bi-nucleated cells and nuclei with areas less than 75 μm^2^, which were likely to be micro-nuclei, were excluded. The migration percentage across the membrane was then calculated as the ratio of the number of cells on the bottom of the membrane to the sum of cells on top and bottom of the filter. For double-stranded DNA damage analysis induced by transwell migration, confocal images of the membranes were acquired with a 40x (oil, 1.4 NA) objective in 200 × 200 μm^2^ fields of view (10 locations per condition). The background fluorescence and average fluorescent intensity of the nuclei were determined as described above. For comparison between the cell types, the results were then normalized to the average fluorescent intensity in the WT MEFs. All experiments were conducted a minimum of two times with two replicates for each group per experiment.

### Atomic force microscopy

All AFM measurements were made with cells in DMEM culture medium using a BioScope II with a Nanoscope V controller (Bruker, Santa Barbara, CA) coupled to an inverted fluorescence microscope equipped with a 10× (NA = 0.3) and 20× (NA = 0.8) objective lenses (Carl Zeiss, Thornwood, NY). Sharp probes were pyramidal cantilevers mounted on a triangular cantilever with a length of 200 μm, a nominal tip radius of 20 nm, a tip half angle of 20° (as measured using scanning electron microscopy), and a nominal spring constant of 0.02 N/m (Olympus TR400PSA; Asylum Research, Goleta, CA). Round probes were 10 μm diameter spheres mounted on silicon nitride cantilevers with a nominal spring constant of 0.01 N/m (Novascan Technologies, Ames, IA). The spring constant was calibrated before each experiment using the thermal fluctuations function of the nanoscope which measures the motion of the cantilever in response to thermal noise. AFM measurements were done at a ramp speed of 800 nm/s, which is sufficiently slow such that our measurements are not rate-dependent, and that with the algorithm used (85), elastic modulus is relatively independent of the indentation and load (61). To avoid substrate effects on AFM measurements, the indentation depths were kept between 100-400 nm (61). Measurements for cytoplasmic and cortical stiffness were performed on regions well away from both nucleus and cell edges while measurements for nuclear stiffness were done above the center of the cell nucleus (18, 20). The Young’s modulus for the AFM sharp and round probe measurements was calculated using models described previously (20). At least twenty measurements were made for each experiment.

### Cellular contractile force measurements

To measure cellular contractile force, we used FTTM (86). Acrylamide-based hydrogels were made on glass bottom 6-well plate as previously described (87) and the Young’s modulus of the gels was 8 kPa, as determined by AFM. After coating the hydrogels with fluorescent markers to visualize displacement and bovine collagen (40 μg/mL) to promote cell attachment, we sparsely seeded cells on said hydrogels and waited overnight. We used an epifluorescence microscope equipped with environment control chamber (Leica DMi8, Germany) to take images of the cells and the fluorescent markers on the hydrogels. Based upon these images, displacements made by cells on the hydrogels were calculated using particle image velocimetry (88) and traction was retrieved from the resulting displacement fields using FTTM (86). As a measure of cellular contractile force, we used both strain energy and net contractile moment, of which the former is less sensitive to the variation in cell spreading area than the latter (86).

### Optical tweezers

A laser beam (10 W, 1,064 nm) was tightly focused through a series of Keplerian beam expanders and a 100x Nikon objective (oil, 1.45 NA). A high-resolution quadrant detector was used for position detection. To measure the mechanical properties of living cells, latex beads with a diameter of 0.5 μm (L3280, Sigma) were added into the culture medium and were endocytosed by the cells overnight at 37°C with 5% CO_2_. To measure the stiffness of the cytoplasm, we chose particles away from both the thin lamellar region at the cell periphery and the nucleus to avoid any interactions with the mechanically distinct cell cortex and nucleus. The selected particle was then dragged away from the nucleus. To measure the stiffness of the nucleus, we selected particles adjacent to the nucleus, and the particle was dragged towards the nucleus. Particles were dragged at a constant velocity of 1 μm/s by the optical trap and the force-displacement curve was recorded. The slope in the linear range of the force-displacement curve was taken as the local stiffness (89).

### Fluorescence recovery after photobleaching (FRAP)

For FRAP studies, cells were seeded into glass bottom 12 well plates (Cellvis, CA) at 100,000 cells/well and transfected with the cDNA constructs encoding EGFP-tagged proteins the next day using Lipofectamine 3000. FRAP studies were conducted 24 hrs post transfection. We used a Nikon A1R confocal microscope equipped with a 30-mW argon 488-nm laser and a 40× (oil, 1.3 NA Plan-Apochromat) objective to perform the FRAP experiments as previously described (42). The region of interest was selected and photobleached for 12 iterations at 20% laser power and the fluorescent recovery was monitored in 2 second intervals at 1% laser power for 360 s. Nikon Elements (NIS-Elements) was then used to quantify the average fluorescence intensity in the region of interest and total cellular intensity. It was then normalized to the changes in total fluorescence intensity as I_rel_ = T_0_I_t_/T_t_l_0_, where T_0_ is the total cellular intensity during the prebleach, T_t_ is the total cellular intensity at time point t, I_0_ is the average intensity in the bleached area during prebleach, and I_t_ is the average intensity in the region of interest at time point t (90). The normalized fluorescence was then plotted against the time after bleaching. To account for the differences of the immobile fraction (the difference between the fluorescence intensity in the bleached area prebleach and the intensity at infinity after bleach) between different cell types and EGFP-tagged constructs, we used a modified time of half-recovery value (t_1/2_) where t_1/2_ is the time after bleach required for the fluorescence levels to reach the median between levels immediately after bleach and prebleach, rather than using the median between prebleach levels and steady-state levels (42). To determine t_1/2_, we used a modification of the method described by Harrington et al. (91). We plotted ln (1 − i_t_) versus time after bleach, where it is the mean normalized fluorescence intensity in the bleach region at time t and 1 is the mean normalized fluorescence intensity in the bleach region prebleach. The curves were fitted using Excel, and t_1/2_ was calculated as t_1/2_ = ln2 × (−1/slope) (42). Data from the first 31 s after bleach were used in all experiments. For each experiment, at least 10 cells were examined, and the t_1/2_ of recovery was calculated from two independent experiments.

### Wound healing assay

Wound healing assays were performed in 35mm μ-dishes with 4 well silicon inserts (Ibidi GmbH). Cells were seeded at 50,000 cells/well and incubated overnight at 37°C with 5% CO_2_. The next day, the inserts were removed, and the migrating cells were imaged for 8 hrs of wound closure time using phase contrast microscopy. Image analysis was done using ImageJ in order to calculate the remaining wound area after 8 hours and the wound closure rate.

### Statistical Analyses

Data are presented as violin plots, box plots, and as mean values ± standard errors (SE). Each experiment was performed a minimum of two times unless otherwise stated. Whole cell and nuclear morphometric data from cell and nuclear morphometrics analyses, AFM, and OT were normally distributed when transformed logarithmically thus, the statistics for these data sets were performed on log-transformed data. Data obtained for fluorescence intensity and cell migration experiments were normally distributed. For all these data sets, ANOVA was used for comparison of the means followed by the Tukey HSD test for multiple comparisons. The distribution of the TFM data was not normal in either the original or log-transformed form. As a result, a Kruskal–Wallis test was used for comparison of the means followed by a Steel-Dwass test for multiple comparisons.

The unpaired Student’s t-test with two tails at the 95% confidence interval was used to determine statistical significance. Linear regression was used to examine the correlation between cell and nuclear volumes, and nuclear sphericity and cell spreading, and the Tukey HSD test was used for multiple comparisons of the slopes of the regression lines slopes. Denotations: *, p < 0.05; **, p < 0.01; ***, p < 0.001; ns, p > 0.05.

## Acknowledgement

The authors thank Daniel Starr for the nesprin-3 antibody; Kyle Roux for the GFP-KASH4 construct; Arnoud Sonnenberg for EGFP-nesprin-3*α*; David Kirchenbuechler for Imaris training and helpful suggestions for imaging data analysis; Christopher Mauer for the FRAP trainings; and Andrew Stephens for insightful discussions and critical review of the manuscript. This work made use of the SPID facility of Northwestern University’s NUANCE Center, which has received support from the SHyNE Resource (NSF ECCS-2025633), the IIN, and Northwestern’s MRSEC program (NSF DMR-1720139). The imaging analysis with Imaris was performed at the Northwestern University Center for Advanced Microscopy generously supported by NCI CCSG P30 CA060553 awarded to the Robert H Lurie Comprehensive Cancer Center.

## Author contribution

A.V.K., R.D.G, S.A.A, Y.Z., J.R.T., G.W.G.L., and G.G.G. conceptualized the work and contributed to the experiments design; S.S. performed the cell cycle analysis studies; F.A.N.S. and S.A.A. created all the knockdown and rescued MEF lines and conducted immunoblotting studies and analysis; Y.L.H. and M.G. conducted optical tweezer studies and analysis; C.Y.P and J.J.F contributed to the traction force microscopy studies and analysis; X.W. and K.L.R. designed and made the construct for the LC knockdown; A.V.K. designed and performed all AFM, immunofluorescence, FRAP, RNAi, and migration studies and analyzed the data; A.V.K., S.A.A, and R.D.G. wrote the initial draft of the manuscript and all co-authors reviewed and edited it.

## Data and materials availability

All of the data that is required for the conclusions is included in the manuscript and/or the Supplementary Material. Any additional data related to this paper may be requested from the authors.

---

**Figure S1:**
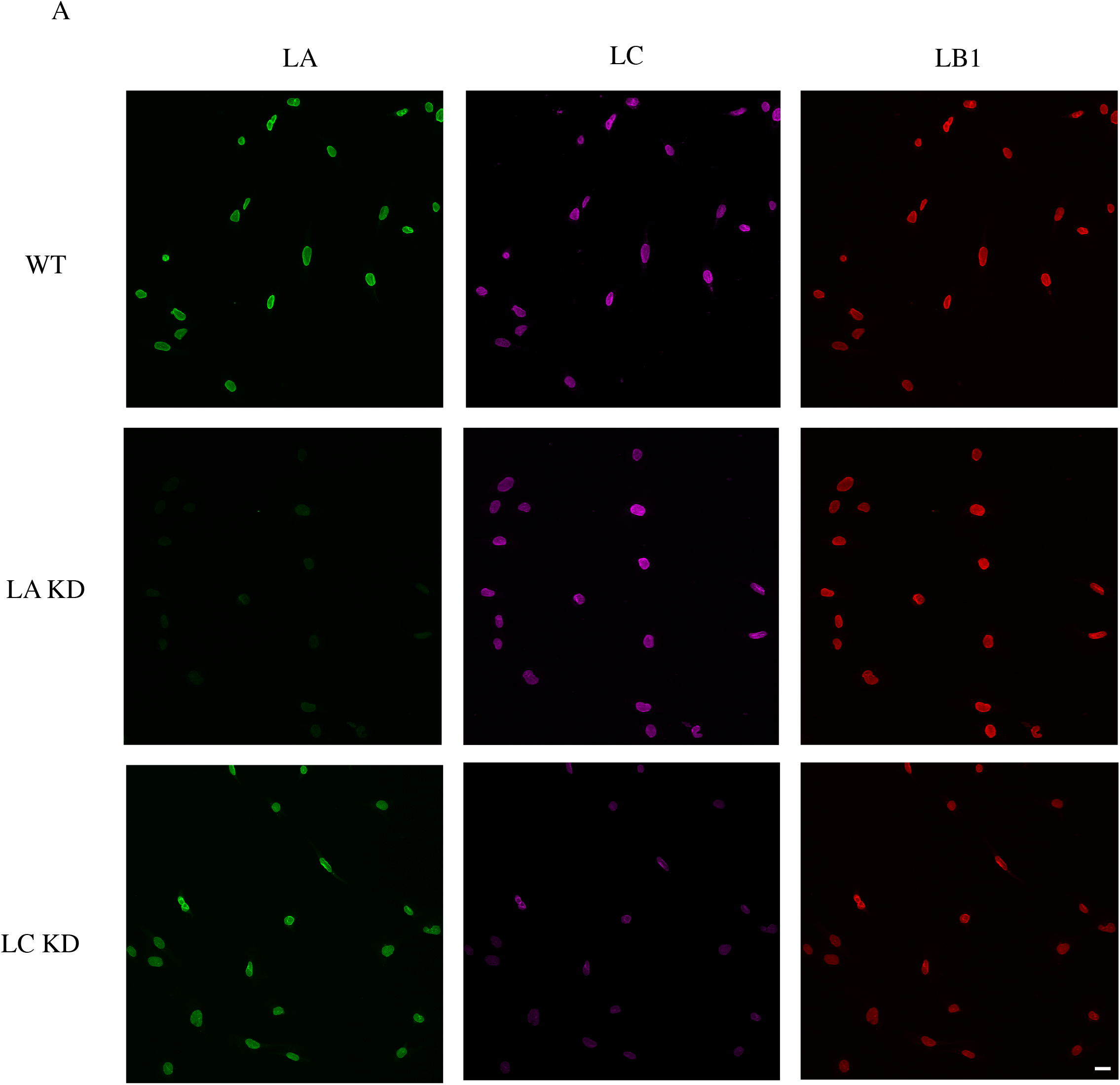
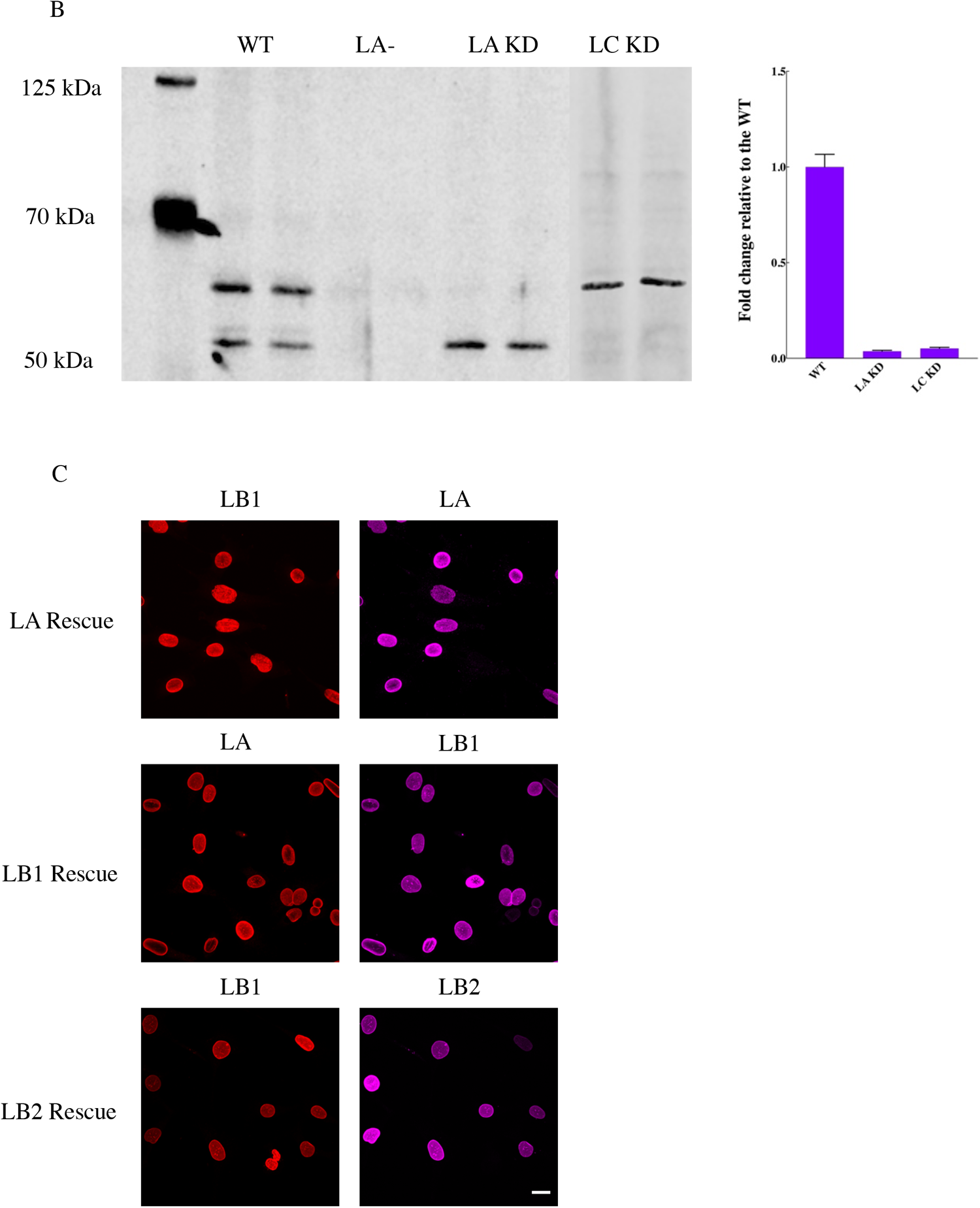
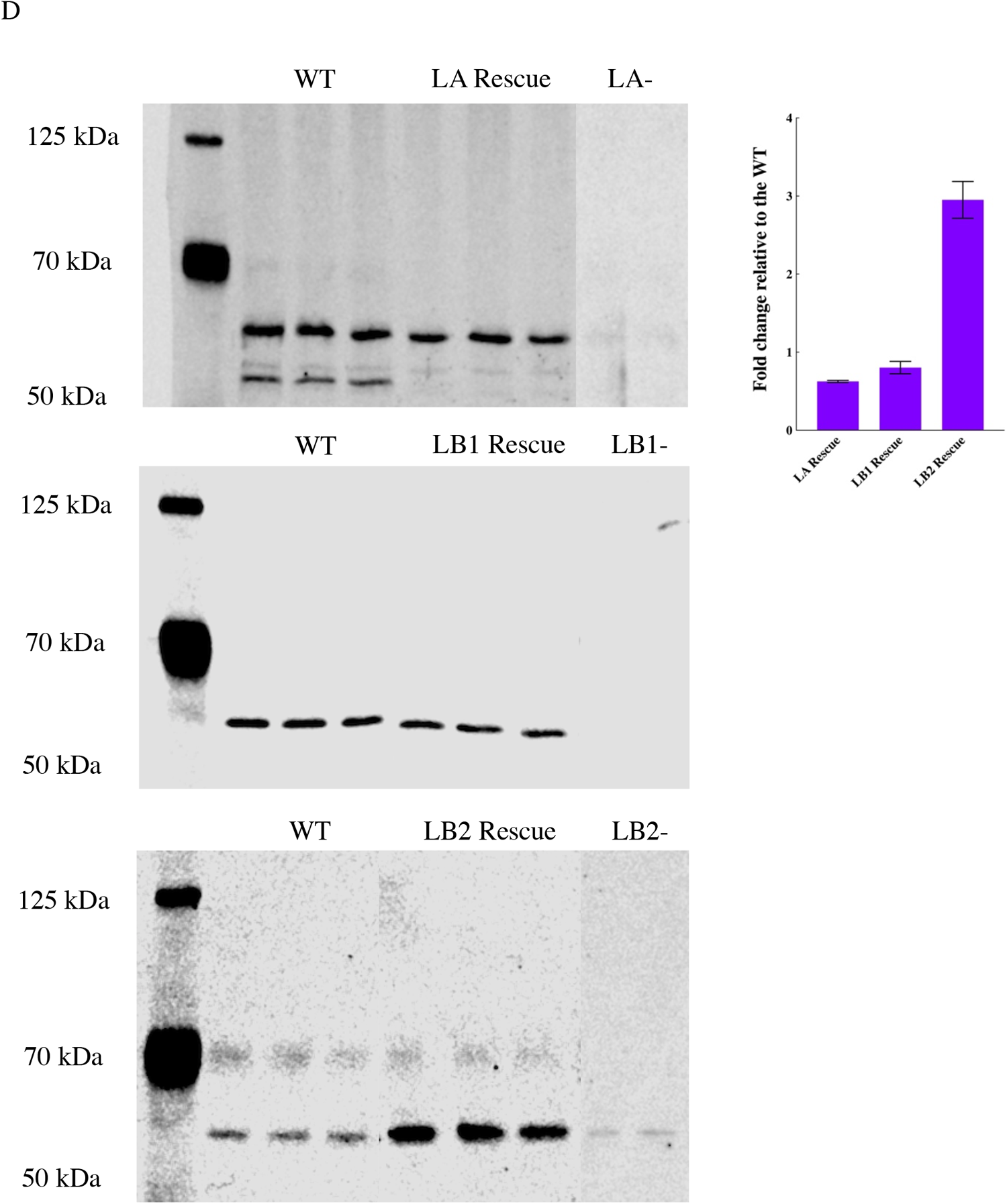
Validation of the lamin knockdown and rescued MEF lines used in this work. **(A)** Representative wide-field immunofluorescence images of the nuclei from the indicated MEF lines stained for anti-LA, anti-LC, and anti-LB1. **(B)** Quantitative immunoblot for the expression level of the knockdown protein in LA KD and LC KD MEFs. **(C)** Representative wide-field immunofluorescence images of the nuclei in LA/C-, LB1-, and LB2-MEFs respectively expressing LA, LB1, or LB2. **(D)** Quantitative immunoblot for the expression level of the rescued protein in LA/C-, LB1-, and LB2-MEFs expressing LA, LB1, and LB2 respectively. Scale bar is 20 μm.

**Figure S2.**
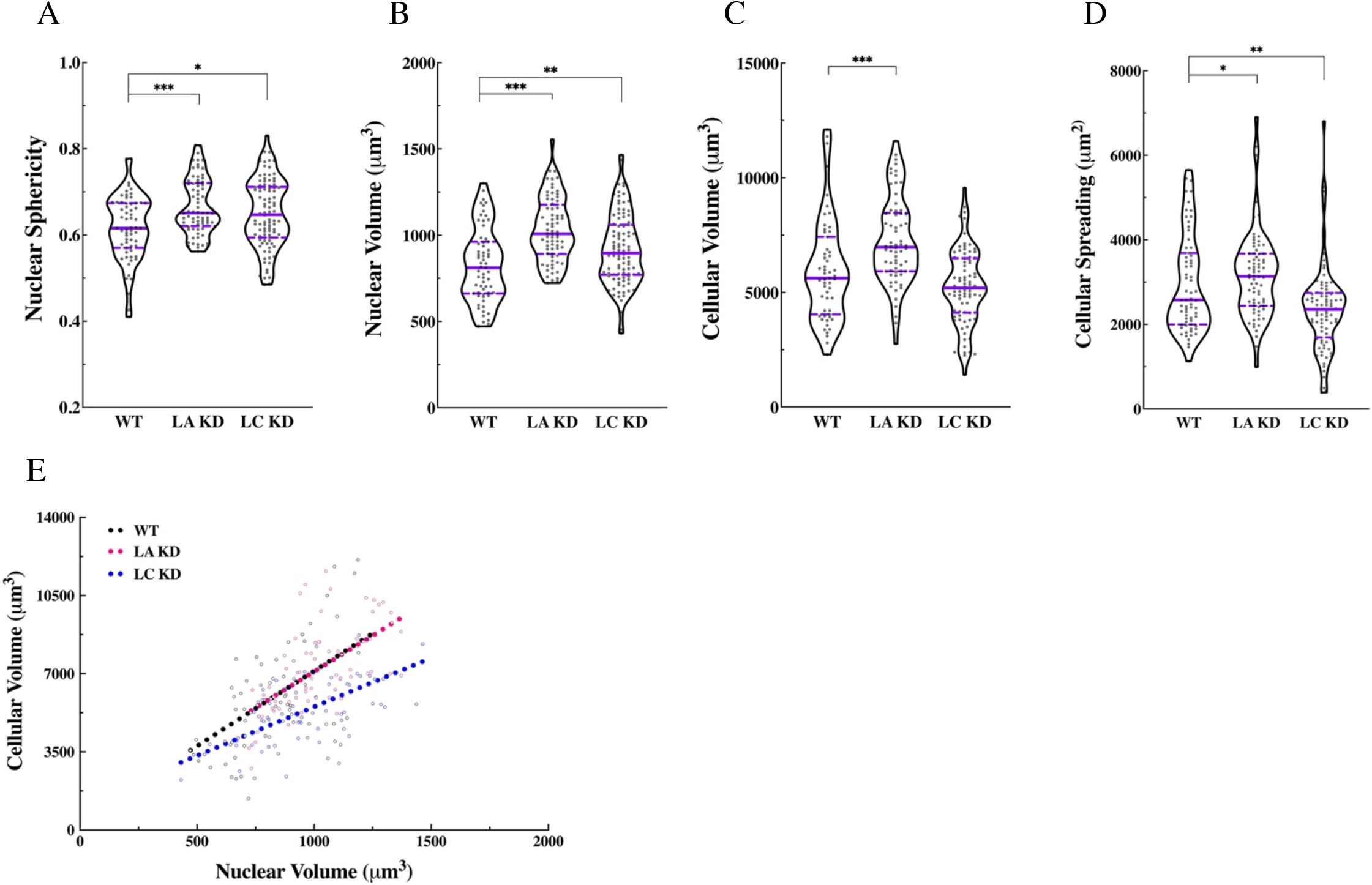
Specific knockdown of LA or LC affects nuclear and cell morphology. Vilolin plots of **(A)** nuclear sphericity, **(B)** nuclear volume, **(C)** cellular volume, **(D)** and cellular spreading area in the WT (n = 3, 55 cells), LA KD (n = 3, 66 cells), and LC KD (n = 3, 82 cells) MEFs. **(E)** Scatter plot of cellular vs. nuclear volume showing the corelation between the two variables in WT (slope = 6.72, *P <* 0.0001), LA KD (slope = 6.47, *P <* 0.0001), LC KD (slope = 4.37, *P <* 0.0001) MEFs. The regression slope for LC KD MEFs is significnatly different than those in WT and LA KD MEFs (*P* < 0.0001). The solid bars in the violon plots represent the median and the dashed lines mark the 25^th^ and the 75^th^ percentiles. \**P* < 0.05, \*\**P* < 0.01, \*\*\**P* < 0.001.

**Figure S3.**
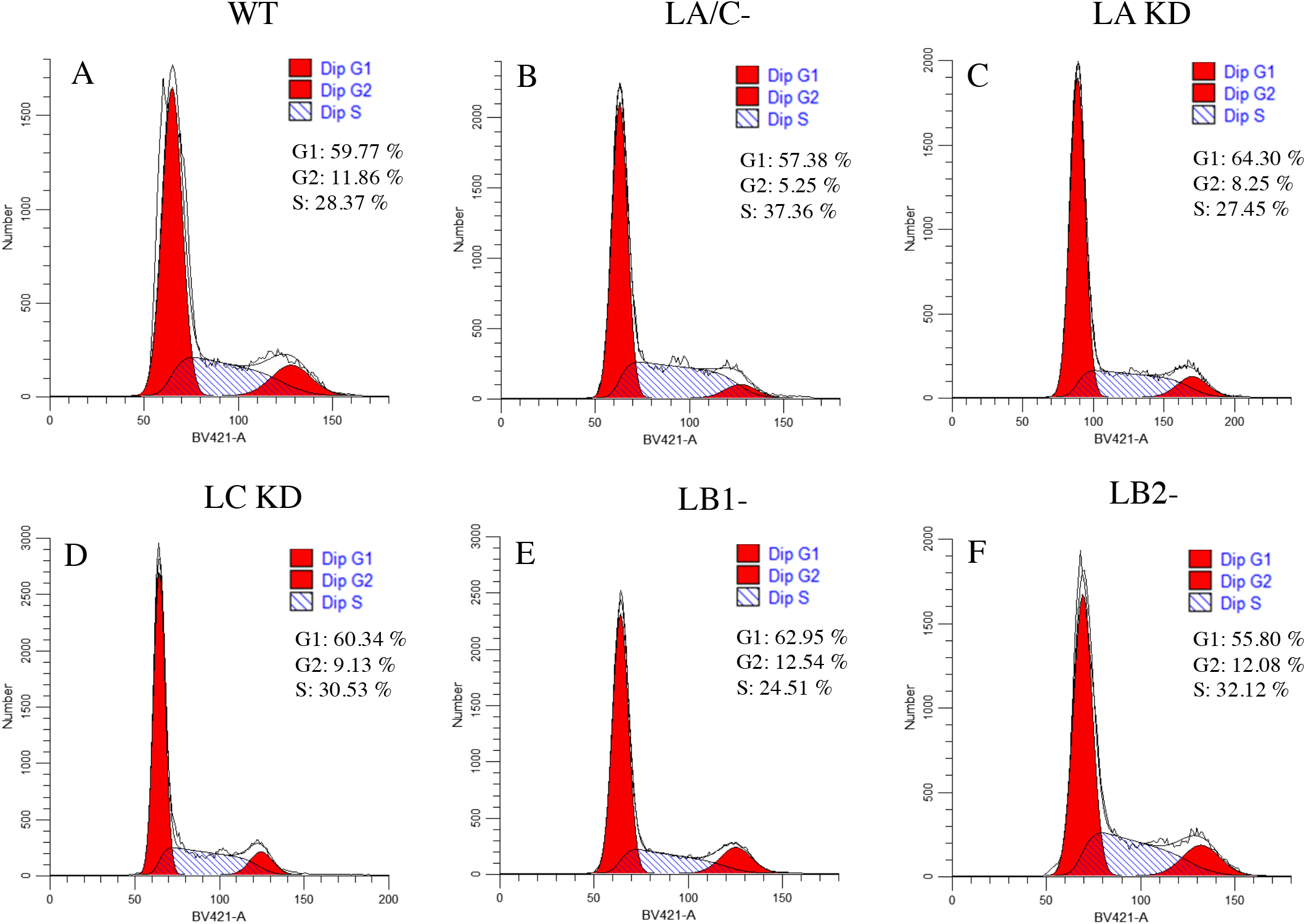
Flow cytometry analysis of the MEF lines used in this work. Histograms of the cell cycle distributions of **(A)** WT, **(B)** LA/C-, **(C)** LA KD, **(D)** LC KD, **(E)** LB1-, and **(F)** LB2-MEFs. The data are quantified as the percentage of the cells in G1, S, and G2 stages of the cell cycle (n=3). The *x* and *y* axes denote the DNA content and cell number, respectively.

**Figure S4.**
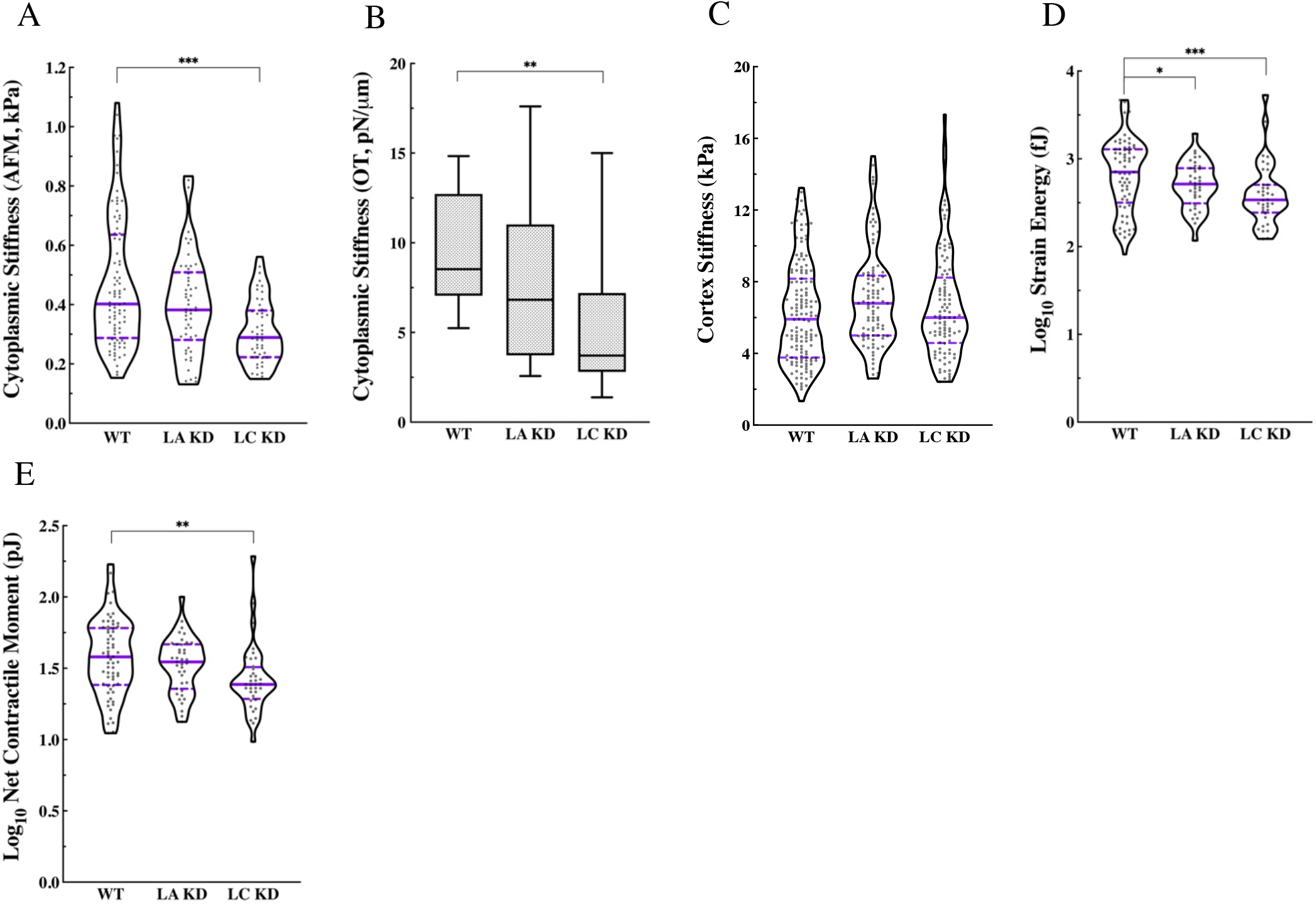
Knockdown of LA or LC compromises cellular mechanics. **(A)** Violin plots of the AFM round probe measurements for the cytoplasmic stiffness in the WT (n = 4, 75 cells), LA KD (n = 2, 55 cell), and LC KD (n = 2, 53 cells) MEFs. **(B)** Box plots of OT measurements of the cytoplasmic stiffness in the WT (n = 1, 9 cells), LA KD (n = 1, 8 cells), and LC KD (n = 1, 13 cells) MEFs. **(C)** Violin plots of the AFM sharp probe measurements for the apical cortex stiffness in the WT (n = 4, 147 cells), LA KD (n = 2, 99 cells), LC KD (n = 2, 100 cells) MEFs. **(D)** Violin polts of the logarithmically transformed SE in the WT (n = 2, 105 cells), LA KD (n = 1, 36 cell), and LC KD (n = 1, 39 cells) MEFs. **(E)** Violin plots of the net contractile moment in the WT (n = 2, 105 cells), LA KD (n = 1, 36 cells), and LC KD (n = 1, 39 cells) MEFs. The solid bars in the violin plots represent the median and the dashed lines mark the 25^th^ and 75^th^ percentiles. The bars and the whiskers in the box plots represent the median and the minimum/maximum, respectively. \**P* < 0.05, \*\**P* < 0.01, \*\*\**P* < 0.001.

**Figure S5.**
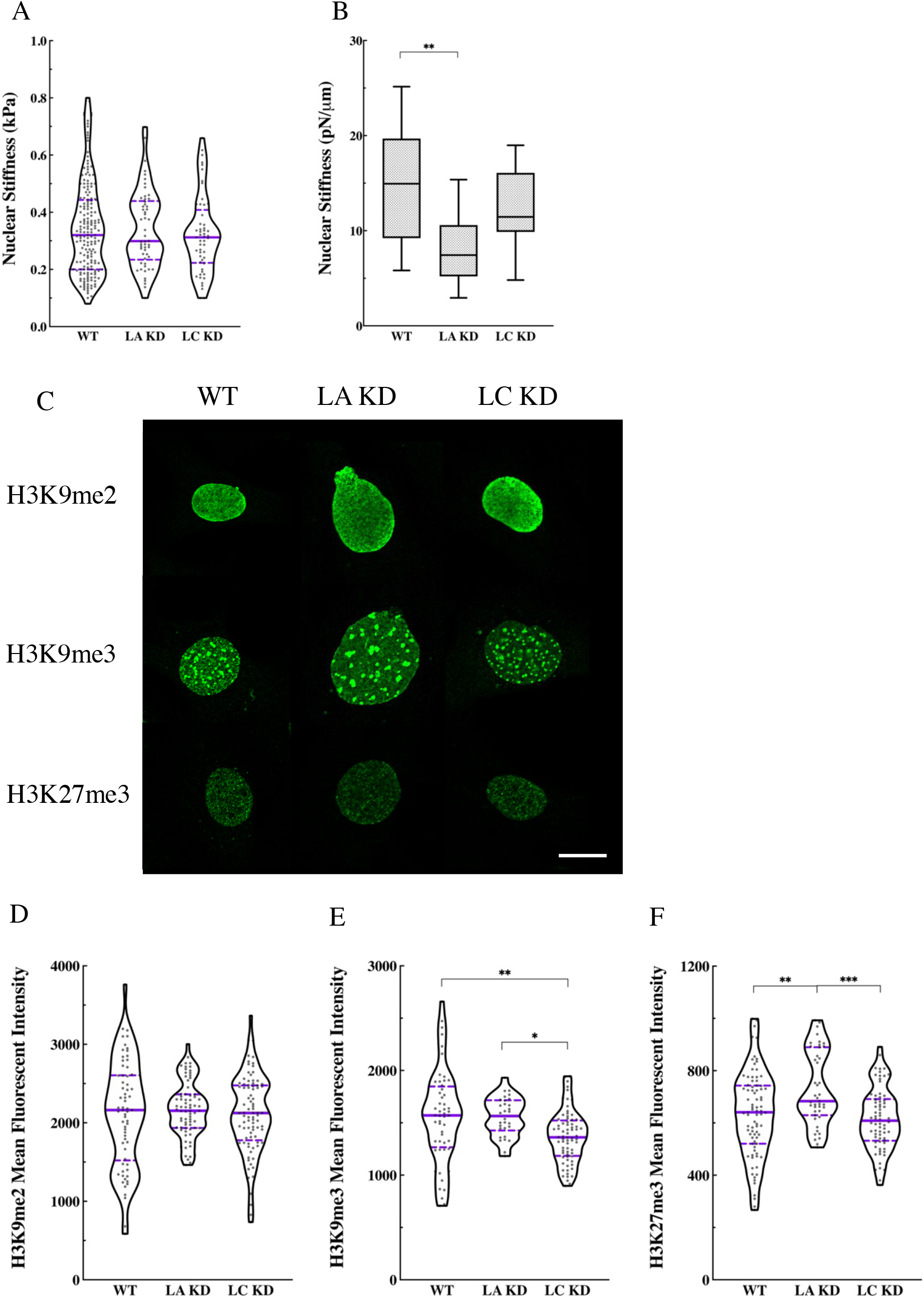
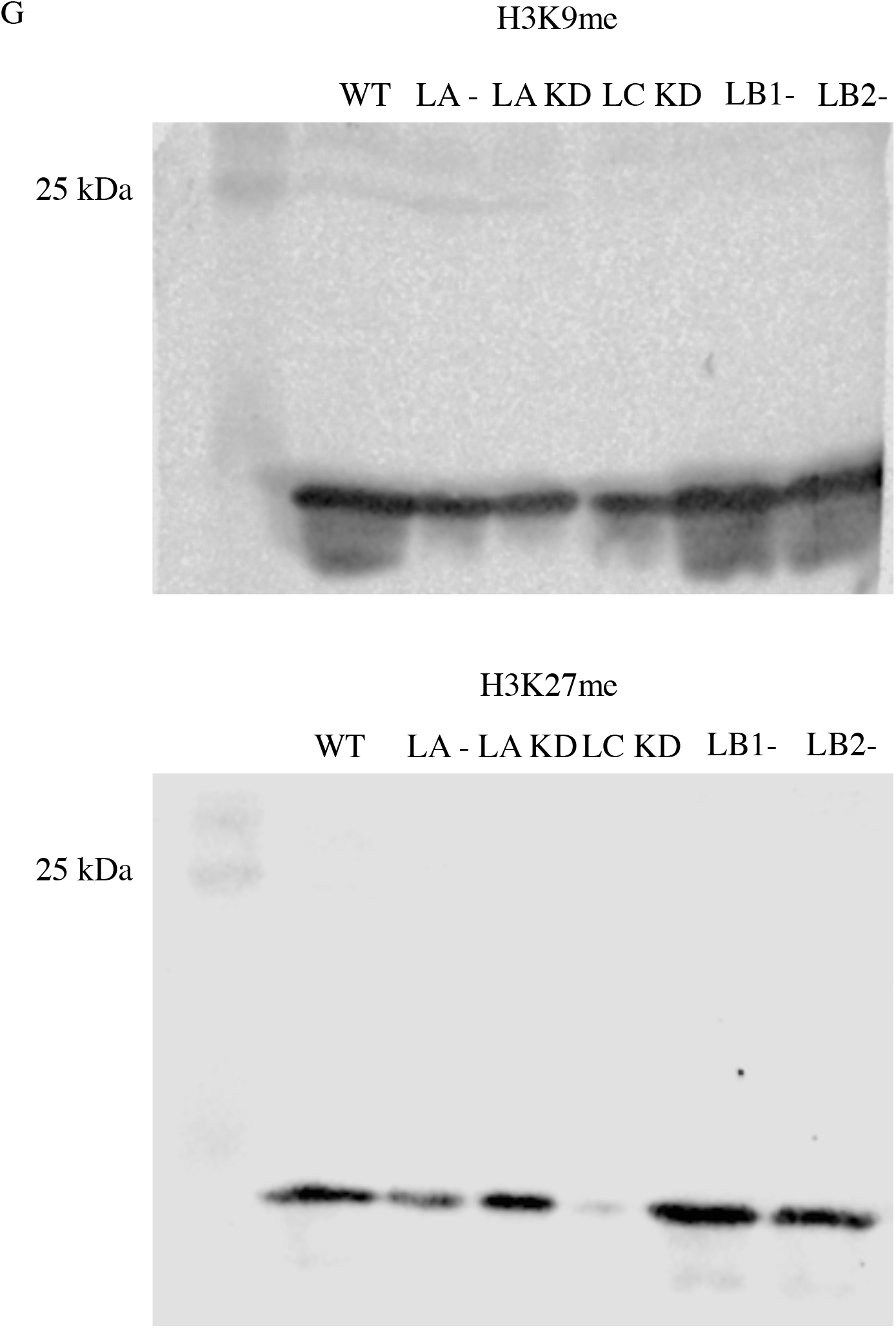
Knockdown of LA or LC affects nuclear stiffness and alters heterochromatin levels in MEFs. **(A)** Violin plots of AFM round probe measurements of the nuclear stiffness in the WT (n = 4, 188 cells), LA KD (n = 2, 54 cell), and LC KD (n = 2, 55 cells) MEFs. **(B)** Box plots of OT measurements for the nuclear stiffness in the WT (n = 1, 9 cells), LA KD (n = 1, 14 cells), and LC KD (n = 1, 9 cells) MEFs. **(C)** Representative maximum projections of the confocal z-stacks of nuclei for the WT, LA KD, LC KD MEFs stained for H3K9me2, H3K9me3, or H3K27me3. Violin plots of the mean fluorescent intensity of **(D)** H3K9me2, **(E)** H3K9me3, or **(F)** H3K27me3 in the WT (n = 2, 54-93 cells), LA KD (n = 2, 30-77 cells), and LC KD (n = 2, 71-85 cells) MEFs. **(G)** Immunoblot for the expression levels of the H3K9me (top) and H3K27me (bottom) in the WT, lamin KO and lamin KD MEFs. The solid bars in the violin plots represent the median and the dashed lines mark the 25^th^ and 75^th^ percentiles. The bars and the whiskers in the box plots represent the median and the minimum/maximum, respectively. \**P* < 0.05, \*\**P* < 0.01, \*\*\**P* < 0.001. Scale bar is 20 μm.

**Figure S6.**
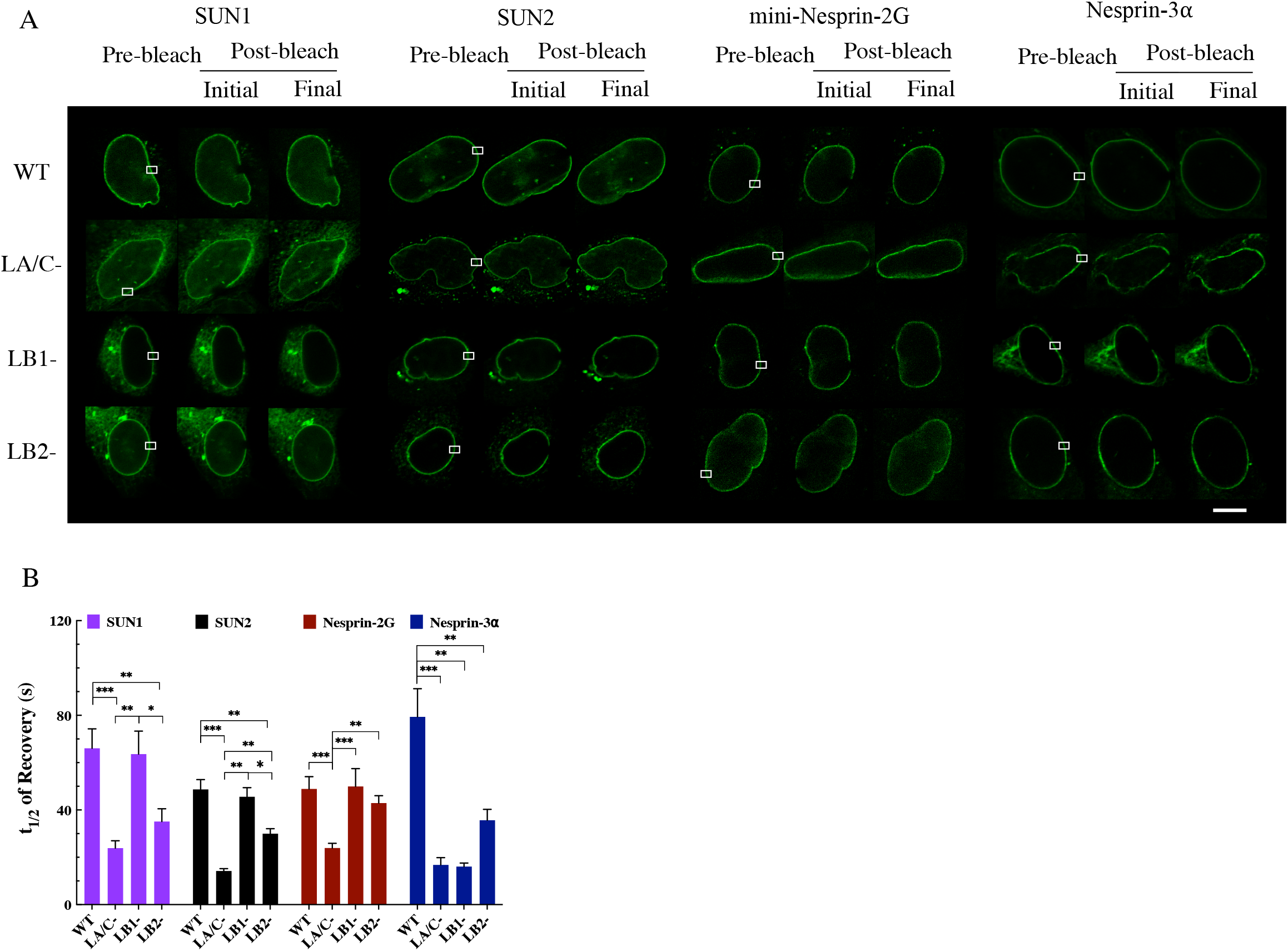
The nuclear lamin isoforms distinctively regulate the dynamics of LINC complex components at the NE of MEFs. **(A)** Representative fluorescense images of the WT, LA/C-, LB1-, and LB2-MEFs lines expressing the indicated EGFP-tagged LINC complex component constructs and subjected to FRAP. The white boxes indicate the photobleached regions of interest. **(B)** Bar plots of the average t_1/2_ of recovery of the indicated EGFP-tagged LINC complex constructs expressed in the WT, LA/C-, LB1-, and LB2-MEF lines (n ≥ 2; 10-15 cells per experimnetal condition). The data are shown as mean ± SE. \**P* < 0.05, \*\**P* < 0.01, \*\*\**P* < 0.001. Scale bar is 10 μm.

**Figure S7.**
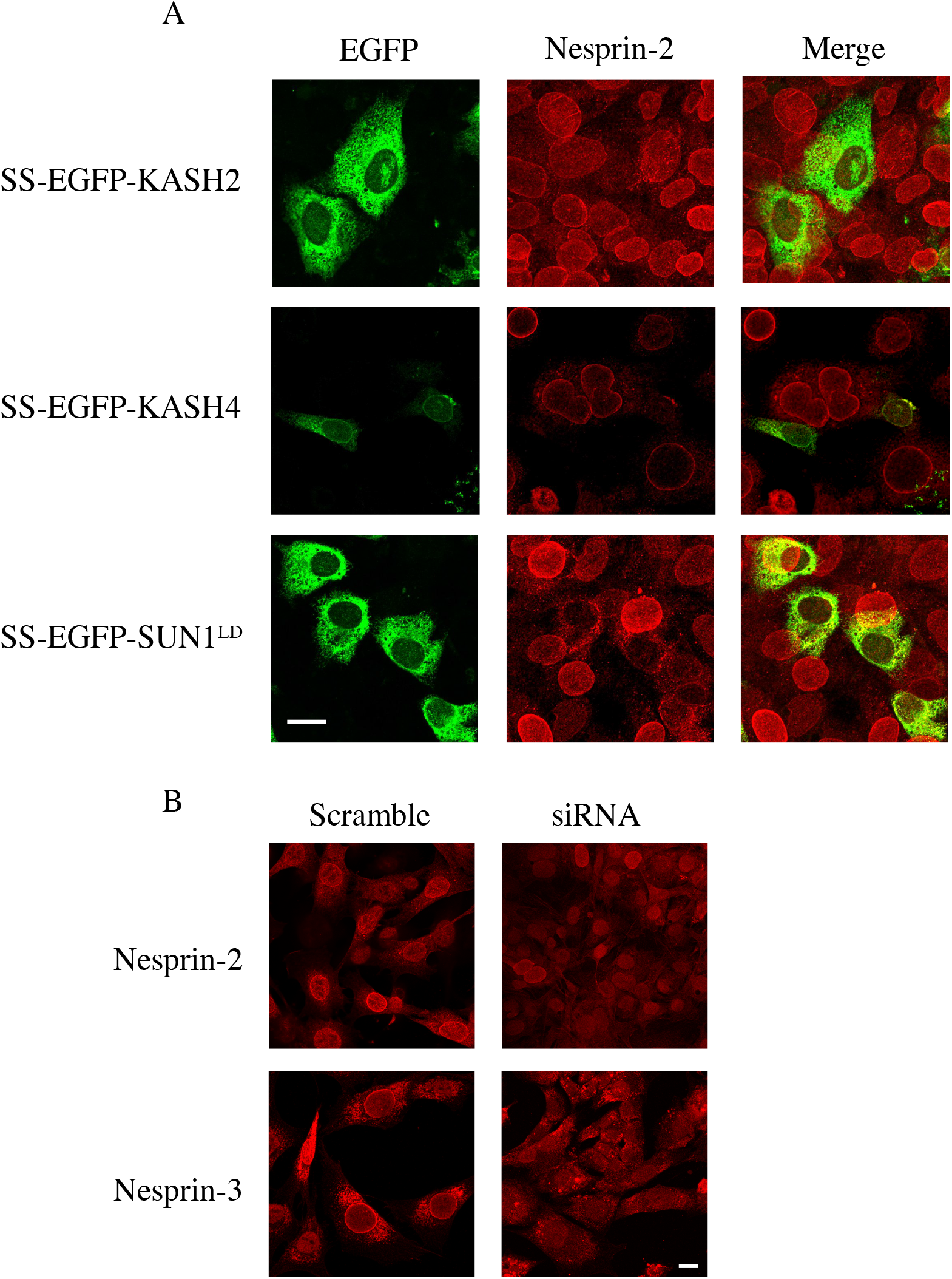
Microscopic verification of the dominant negative inhibition or selective siRNA disruption of LINC complexes. **(A)** Representative wide-field immunofluorescence images of nesprin-2G staining in the WT MEFs expressing the indicated SS-EGFP-tagged constructs. **(B)** Representative wide-field immunofluorescence images of nesprin-2G (top) or nesprin-3α (bottom) staining in a MEF line (shown here is LB2-) treated with the nesprin-2G (top) or the nesprin-3α (bottom) targeting siRNA or the non-coding control siRNA followed by staining for nesprin-2 (top) and nesprin-3 (bottom). Scale bar is 20 μm.

**Figure S8.**
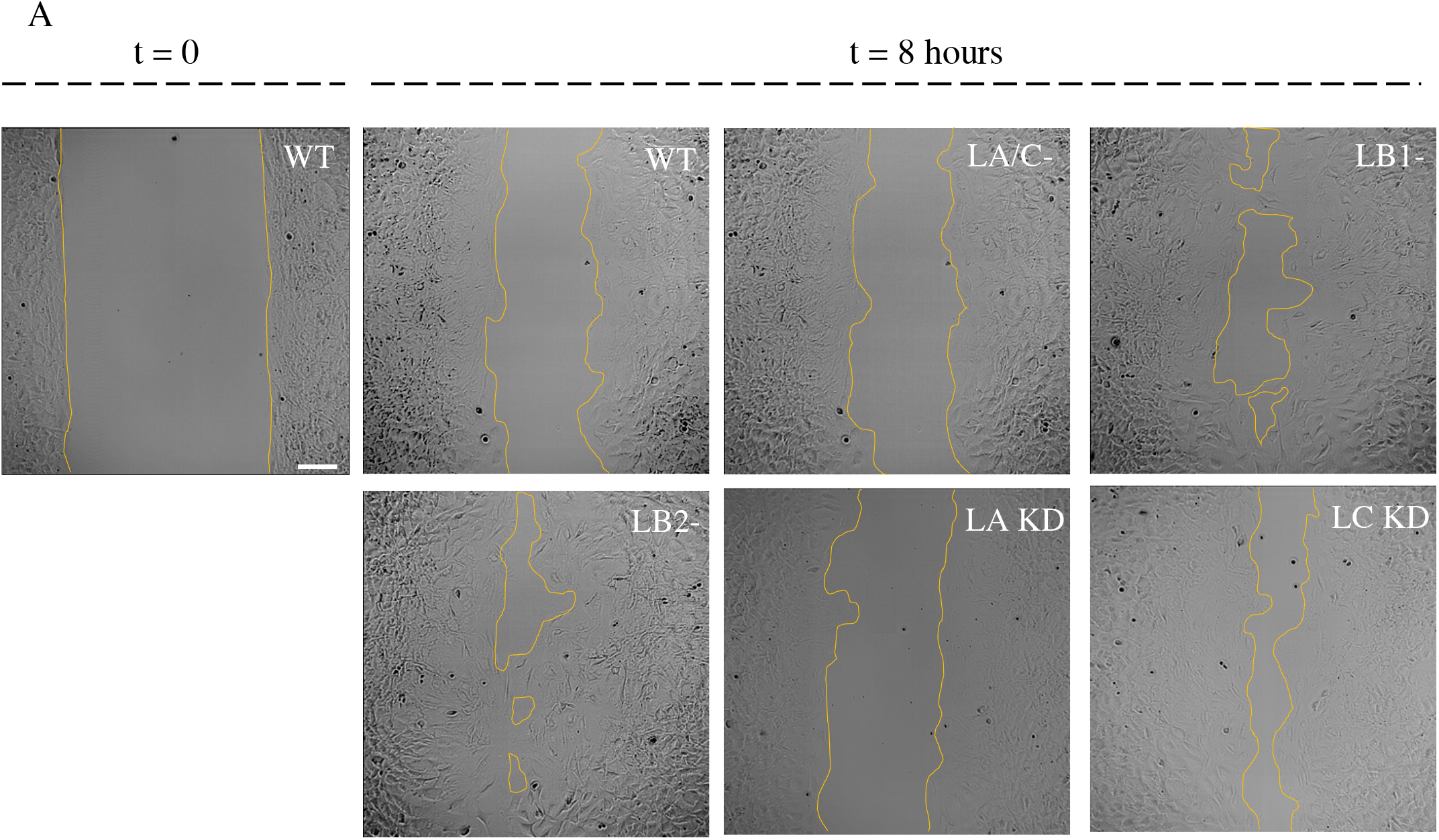
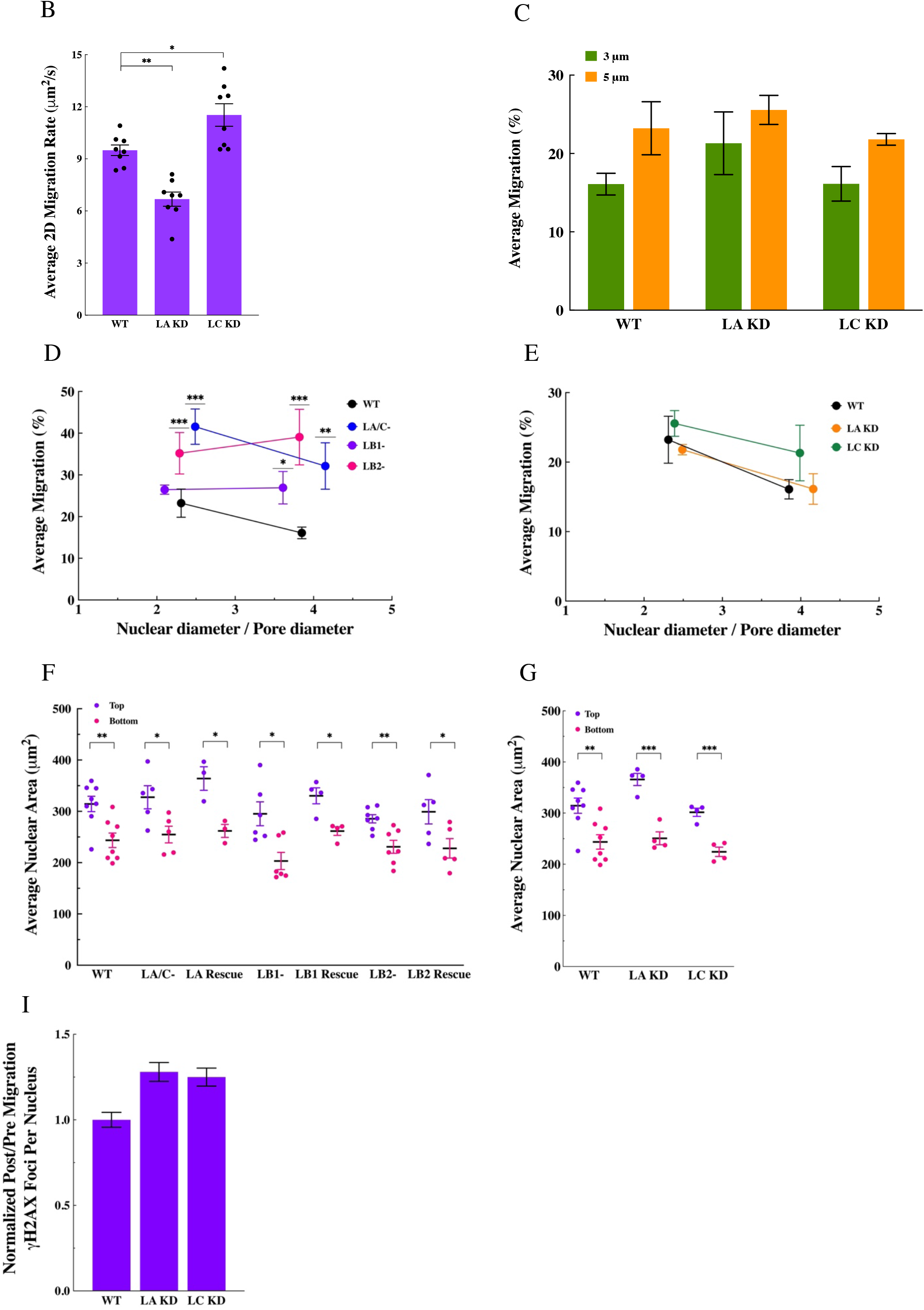
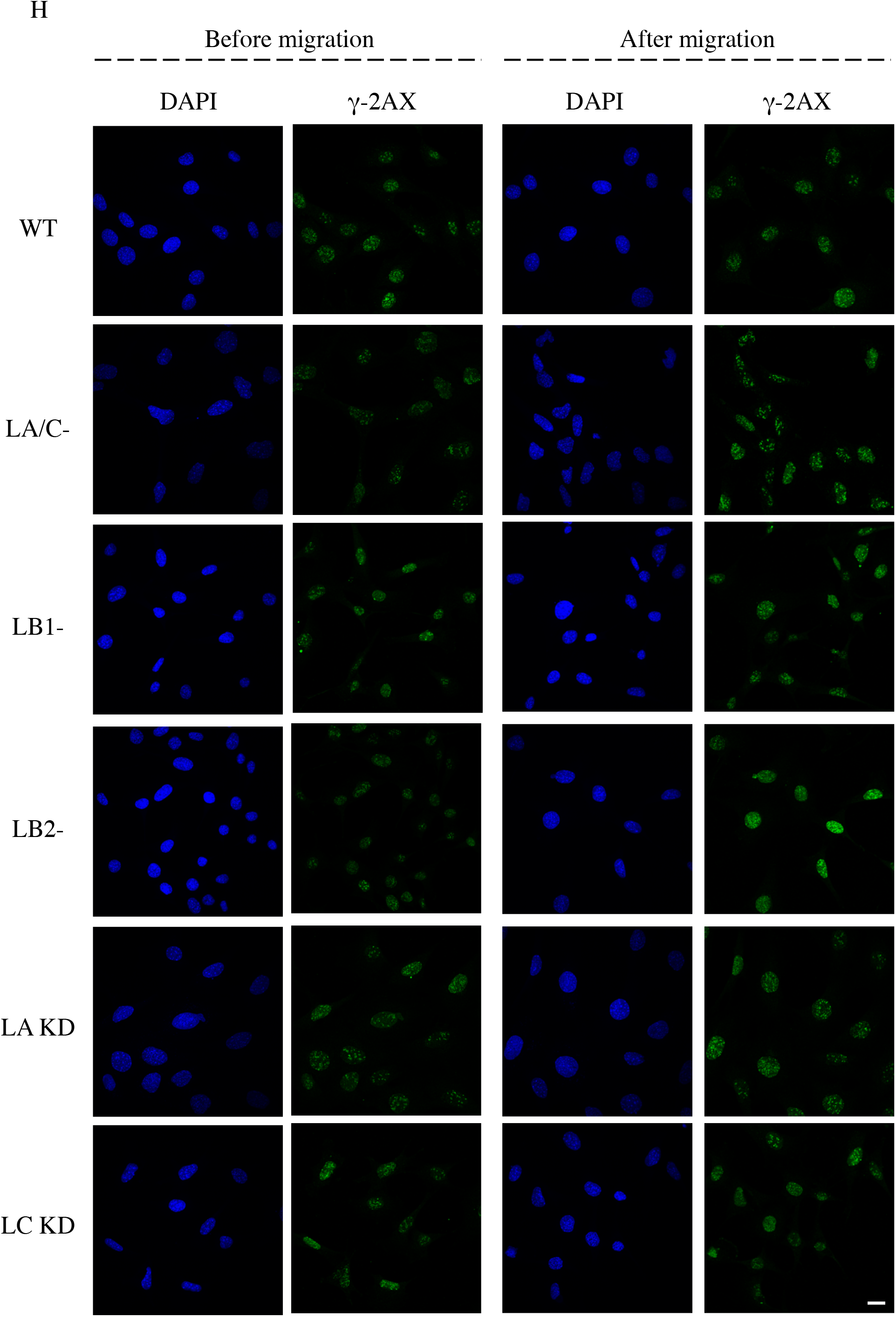
Loss or knockdown of lamin isoforms changes cell migratory behavior and migration induced DNA damage. **(A)** Representative phase contrast images of the indicated MEF lines migrating in a 2D wound healing assay for 8 hours. The t = 0 is when the silicon insert is removed from the dish. The yellow lines demarkate the leading edge of the wound. Scale bar is 100 μm. **(B)** Bar plots of the average 2D migration rate during the 2D wound healing assay for the WT, LA KD, and LC KD MEFs (n = 8). **(C)** Bar plots of the average migration percentage for the WT, LA KD, and LC KD MEFs challenged with transwell membranes with 3 μm or 5 μm pore diameters (n ≥ 3 for each set of experiments). Comparison of the nuclear to pore diameter ratio and migration through 3 μm and 5 μm pores in **(D)** lamin KO and **(E)** lamin KD MEFs to the WT MEFs (n ≥ 50 cells per condition). **(F)** Scatter plots of the average nuclear area for the indicated MEF lines present at the top and bottom of the transwell membranes. **(G)** Same as **(F)** for the the WT, LA KD, and LC KD MEFs. (n ≥ 3, at least 50 cells per experiment condition). **(H)** Representative maximum projections of confocal z-stacks of DAPI- and γ-H2AX-stained nuclei in the indiated MEF lines before and after their migartion through transwells with 3 μm pores. Scale bar is 20 μm. **(I)** Bar plots of the normalized post/pre migration γ-H2AX foci counts in the WT, LA KD, and LC KD MEFs that migrated through transwell membranes with 3 μm pores. (n = 3, at least 150 cells per experiment condition). Data are shown as mean ± SE. \**P* < 0.05, \*\**P* < 0.01, \*\*\**P* < 0.001.

